# Craniofacial suture complexity and digging mode importance in rodents

**DOI:** 10.64898/2026.09.11.750232

**Authors:** Sarah Saxton Strassberg, Kenneth D. Angielczyk

## Abstract

The morphology of cranial sutures (i.e., joints that connect the bones of the skull, are sites of growth, and absorb biomechanical stresses) is highly variable, especially in mammals. Exposure to different strain regimes as a result of development, function, and ecology can drive disparity in suture complexity at both the individual and taxonomic levels. Although the relationships between diet, muscle mass, static loading, and certain behaviors (e.g., headbutting) and suture morphology are well studied, the effects of fossoriality, particularly the use of the craniodental apparatus during substrate excavation, remain largely unknown. We synthesize data using an ordinal ranking scheme for digging mode importance in 107 rodent species; premaxillofrontal and nasofrontal suture complexity as measured by sinuosity index, power spectrum density, and spectral entropy, the latter of which is a novel suture complexity metric; and Bayesian multilevel models to examine the fossoriality-suture phenotype relationship at the suborder level and above and predict digging mode usage in 11 cryptic extant species. Most fossorial rodents utilize multiple digging modes, often at varying levels of importance. Chisel-tooth digging importance is associated with reduced nasofrontal suture complexity in multiple suborders, whereas scratch digging importance is correlated with elevated complexity in the premaxillofrontal suture. Suture complexity poorly reflects head-lift digging importance. We demonstrate that fossorial function is a complex, multivariate trait and reveal new insights into the functional morphology of cranial sutures across the most diverse mammalian order.

## Introduction

### Cranial sutures as reflectors of function and ecology

Cranial sutures are connective tissue articulations that join the bones of the vertebrate skull (Moss 1957, Opperman 2000, Di Ieva et al. 2013). Sutures act as “strain sinks,” preventing cranial bones from experiencing high stresses (Rafferty and Herring 1999, Rafferty et al. 2003, Sharp et al. 2023), and experience prolonged bone deposition and growth during ontogeny (Moss 1957, Miura et al. 2009, Gorucu-Coskuner et al. 2023). Particularly within Mammalia, ectocranial suture morphology is highly variable between sutures (e.g., sagittal, nasofrontal), ontogenetic stages, and taxa, ranging from simple and linear to remarkably interdigitated and complex (Buezas et al. 2017, White et al. 2020, White et al. 2025).

Although general suture structure is hereditarily determined, manifestation of specific morphological qualities largely depends on environmental factors (Oudhof 1982). Biomechanical stresses are a substantial driver of cranial suture morphological variability. Exposure to tensile forces tends to result in simple sutures with shallow interdigitations, whereas compressive forces drive complex, deeply interdigitated morphology (Rafferty and Herring 1999). In turn, suture complexity has adaptive significance. Elevated suture sinuosity confers greater bending strength and energy absorption, with the strength of very highly interdigitated sutures approximating that of pure bone (Jaslow 1990, Maloul et al. 2014). Sutures with greater morphological irregularity may also exhibit elevated ductility, damage tolerance, and strain to failure (Liu et al. 2017). At the individual level, sutures have been shown to remodel in response to various *in vivo* experimental strain regime shifts, including jaw or neck musculature removal (Washburn 1947, Moss 1962) or hypertrophy (Byron 2004), changes in dietary hardness (Engstrom et al. 1986, Byron et al. 2018, Byron et al. 2023), and artificial static loading (Hinrichsen and Storey 1968, Wagemans et al 1988). At the species level or above, harder diets in primates (Byron 2009, Byron et al. 2023), larger horns and headbutting behavior in male wild sheep (Jaslow 1989), increased masticatory strain in suoids (Herring 1972), and even cranial growth patterns in humans (Anton et al. 1992, Blumer et al. 2024) correlate with increased suture complexity, indicating taxon-level adaptation.

The effects of locomotion, including fossoriality (i.e., burrowing or otherwise moving earthbound substrate), on suture morphology have received relatively little attention, especially at higher taxonomic levels. Excavation of substrate, particularly the usage of digging modes (i.e., skeletal elements and types of motion used to dig) that involve the craniodental apparatus (see Nevo 1979, Hildebrand 1985, Lacey 2000), likely place elevated biomechanical stresses on the skull and therefore may affect suture morphology at the species level and above. For example, in multiple amphisbaenian lineages, the most complex cranial sutures are oriented transversely to resist stresses induced by headfirst burrowing (Gans 1960, Gans 1978). Within Caviomorpha (Rodentia: Hystricomorpha), the two clades that most frequently engage in chisel-tooth digging (Ctenomyidae and Octodontidae) exhibit elevated suture complexity compared to other families (Buezas et al. 2017). In the fossil record, burrowing dicynodont therapsids have elevated nasofrontal suture complexity, with cistecephalids, the most specialized fossorial clade, exhibiting the most extreme complexity scores (Kammerer 2021).

### Fossorial function in Rodentia

Rodents are an ideal system in which to examine the relationship between suture morphology and fossorial function. Rodentia is the most taxonomically and arguably ecologically diverse clade of mammals, with over 2,000 extant species and ecomorphotypes ranging from gliding to ricochetal, semiaquatic, and subterranean (Auffray et al. 2009, Smiley et al. 2020). Habitual fossoriality has evolved in numerous rodent lineages, with multiple groups converging on highly subterranean lifestyles (e.g., Spalacidae, Geomyidae, Bathyergidae) and others exhibiting lower degrees of specialization for digging (e.g., extant Castoridae, many Sciuridae) (Nevo 1979, Whitford and Kay 1999, Lacey 2000, Hopkins 2005, Bhagat et al. 2021). Unlike other mammalian groups, which tend to be limited to usage of a single digging mode, many fossorial rodent lineages can utilize multiple digging modes. Three primary digging modes are observed in extant rodents: scratch (i.e., quick, running-like motions of the forelimbs), chisel-tooth (i.e., usage of the incisors), and head-lift (i.e., usage of the skull/snout as a wedge) (Lacey 2000, Marcy et al. 2013, Gomes Rodrigues et al. 2023). A given species may use multiple digging modes interchangeably or at different frequencies, often in response to environmental factors. For instance, the naked mole-rat (*Heterocephalus glaber*) digs primarily with its incisors but utilizes its claws and skull secondarily (Tucker 1981, Lacey 2000), and many tuco-tuco (Ctenomyidae) species will favor usage of either their claws or incisors depending on soil hardness (Kubiak et al. 2018, Vasallo et al. 2021).

Chisel-tooth and head-lift digging have evolved multiple times across Rodentia, giving rise to remarkable morphological, functional, and behavioral convergence (Hildebrand 1985, Lacey 2000, Marcy et al. 2016, Gomes Rodrigues et al. 2023). However, relatively few studies have examined functional morphology associated with multiple digging modes in tandem, and most of these focus on a single lineage (Marcy et al. 2016, Morgan et al. 2017, Kubiak et al. 2018, Vasallo et al. 2021). Incorporation of information regarding frequency of usage of a given digging mode or its importance to a species’ overall lifestyle is rare in comparative studies. However, data pertaining to digging mode importance are readily available for numerous taxa (e.g., Eisenberg 1975, Butynski and Mattingly 1979, Tucker 1981, Nowak 1999, Marcy et al. 2013, Vasallo et al. 2021), allowing for more nuanced examination of fossorial function. Behavior and ecology exist on spectra and may differ even between conspecifics to an extent. However, an ordinal ranking scheme (see Wisniewski et al. 2023) for digging mode importance can facilitate more comprehensive understanding of fossorial locomotion and ecology than traditional binary or qualitative categorization methods and provide novel insights into the form-function relationship and ecological niche structure. Additionally, many extant species are cryptic and lack substantial published behavioral, functional, or ecological data. Using phenotype to predict function in such taxa is crucial for better understanding their biology and broader eco-evolutionary patterns.

In this study, we test two hypotheses: 1) that increased specialization for chisel-tooth and head-lift digging, but not scratch digging, correlates with elevated cranial suture complexity in rodents, and 2) that multiple metrics of suture complexity reflect digging mode importance equally strongly across rodent suborders. We collected morphological data, including suture complexity metrics, from 520 specimens representing 107 extant rodent species, and performed a comprehensive literature review to evaluate the importance of scratch, chisel-tooth, and head-lift digging to each species. We fit Bayesian multilevel models with suture complexity metrics as predictors and digging mode importance rankings as ordinal response variables to examine the strength of the relationship between suture morphology and fossorial function across rodent suborders. We then used the models with best performance and greatest strength of support to predict digging mode importance in 11 data-deficient extant species.

## Materials and Methods

### Sampling

520 specimens representing 107 species, all five rodent suborders, and 24 families were sampled. Because suture morphology changes across ontogeny (Gorucu-Koskuner et al. 2023, Khurelbaatar et al. 2025), only adult specimens (i.e., those with a functional third molar) were selected. Specimens were sourced from the Field Museum of Natural History (FMNH) and the Smithsonian National Museum of Natural History (NMNH). Sampled species were chosen to bracket extant phylogenetic and ecological diversity across Rodentia and capture the majority of transitions into head-lift and chisel-tooth digging. To minimize the effects of intraspecific dietary, functional, and environmental variability on suture morphology, specimens for a given species were selected from the same locality and collection timeframe whenever possible, and wild-caught specimens were prioritized. See Supplementary Data for a complete list of specimens used. A phylogeny of the sampled species was derived from Upham et al. (2019); the maximum clade credibility tree was obtained from 10,000 posterior trees.

We selected the premaxillofrontal and nasofrontal sutures as the focal point for this study. Suture morphology in the craniofacial region responds readily to experimental manipulations, including jaw and neck muscle alteration and shifts in dietary hardness (Moss 1957, Herring 1972, Engstrom et al. 1986, Byron et al. 2004). The morphology of these two sutures also varies considerably across taxa, including between fossorial and non-fossorial lineages (Buezas et al. 2017, Kammerer 2021). Craniodental digging modes can subject this region of the skull to elevated biomechanical strains (Gomes-Rodrigues et al. 2023, Vasallo et al. 2021) and thus may affect the associated suture morphology.

Two-dimensional photographs in dorsal view were taken using a Canon Eos Rebel T5i; for skulls with snout-to-occiput lengths of less than 6 cm, a Canon 100 mm f/2.8L Macro IS USM was used. This approach was chosen over using microCT scans or other costly and time-intensive three-dimensional imaging techniques to prioritize overall sample size and taxonomic breadth. Although suture tracings from 2D photographs tend to have slightly higher complexity scores due to parallax, differences between species have been shown to be consistent (Byron 2009), and there is no evidence to suggest differences in suture tracing quality between 3D scans and high-quality 2D photographs (Byron et al. 2023). Additionally, the effects of parallax may be ameliorated by the relatively planar morphology of the dorsal craniofacial region in most rodent taxa sampled in this study; exceptions include the South African springhare (*Pedetes capensis*) and both Old World (Erethizontidae) and New World porcupines (Hystricidae), which tend to have more globular facial regions.

To minimize the effects of asymmetry, only the left nasofrontal and premaxillofrontal sutures were sampled. All measurements and suture tracings were collected in FIJI v2.16.0 (Schindelin et al. 2012). Images were processed at 300% magnification, and scales were set using scale bars present in the photographs. Sutures were traced as freehand lines using a Wacom Intuos Creative Pen Tablet CTL-4100. Landmarks were placed at the endpoints of each suture, and basal skull length was measured as the linear distance from rostrum to occiput. Suture tracings and landmarks were saved as regions of interest (ROIs). To ensure consistency in suture tracing methodology, all tracings and endpoints were digitized by SSS within a three-month period.

ROIs and the associated image TIFF files were imported into RStudio (version 2024.04.2+764) using the RImageJROI package (Sterratt et al. 2014). All subsequent analyses were performed in this version of R. Arc-length parameterized splines were fit to each suture tracing to reduce pixel-level noise while still capturing morphology, and each resulting curve was resampled to have 512 equally spaced semilandmarks. This number was chosen to both adequately capture morphology in even the most complex sutures and to best facilitate subsequent Fourier transforms.

### Digging mode importance ranking scheme

Importance ranks for scratch, chisel-tooth, and head-lift digging were assigned to each species based on published behavioral and ecological data from the primary literature (e.g., ecological, behavioral, or kinematic studies) and comprehensive reference sources (e.g., *Walker’s Mammals of the World*, *Handbook of the Mammals of the World*, *Mammals of the Neotropics*). Literature with primary data or comprehensive synthesis of published behavioral and ecological data was prioritized, and literature with unsubstantiated, anecdotal, or conjectural claims of digging mode usage was given less weight. If the fossorial habits or general ecology of a species were not sufficiently described in the literature and/or if information regarding digging mode usage was lacking, unclear, or strongly conflicting across sources, the species received a ranking of “Unknown.” See Supplementary Table 1 for species rankings and sources used. For data-rich species, the ranking system was established as follows.

*Rank 1*: The digging mode is used extremely rarely or not at all. This rank was typically assigned if no sources described a species utilizing this digging mode and/or at least one source remarked upon a lack of usage. If at least one source described a species using a certain digging mode but the data were anecdotal or conjectural in nature or suggested extreme or unusual circumstances, the species received a rank of 1 for this digging mode. For example, Agrawal (1967) described a number of taxa (e.g., *Marmota, Hystrix*) as using a combination of claws and incisors to dig but did not cite any sources nor provide elaboration on behavioral or functional observations. Because we could not find additional information about the usage of chisel-tooth digging in these taxa, they received a rank of 1 for this digging mode. Likewise, see Burns et al. (1989) for discussion of white-tailed prairie dog (*Cynomys leucurus*) burrows that contained incisor marks but were inferred to be the product of desperation and likely resulted in substantial physical damage to the animals; this species was assigned a rank of 1 for chisel-tooth digging.

*Rank 2*: The digging mode is of marginal importance to the species’ function and lifestyle, used sparingly or only in specific scenarios, such as excavating especially hard or soft soils, removing obstacles (e.g., rocks, roots) from the animal’s path, or moving loosened soil after primary excavation. If a taxon uses all three digging modes but one is utilized at substantially higher rates than the others, the two less commonly used modes were both given ranks of 2. For example, *Heterocephalus glaber* primarily uses its incisors for excavation but occasionally utilizes its claws and snout as well (Jarvis and Sale 1971, Tucker 1981). Therefore, it received a ranking of 2 for scratch and head-lift digging and a ranking of 3 for chisel-tooth digging. Some taxa (e.g., the groundhog, *Marmota monax*; the spinifex hopping mouse, *Notomys alexis*; the plains viscacha, *Lagostomus maximus*) solely use the forelimbs for the primary action of excavation but remove and/or pack loosened soil with their snouts or heads afterwards; such species were assigned a rank of 2 for head-lift digging. Species that scratch dig but exhibit a primary ecological role that is not fossorial in nature (e.g., the pacarana, *Dinomys branickii*; the North American beaver, *Castor canadensis*) were assigned a scratch digging rank of 2 to reflect overall lower degrees of specialization for substrate excavation.

*Rank 3*: The digging mode is of high importance to the species’ ecology and overall lifestyle and is typically used for the primary action of excavation. If a taxon uses more than one digging mode approximately equally and both are of substantial, rather than marginal, importance to its digging-related function and general lifestyle, both were assigned a rank of 3 (e.g., chisel-tooth and head-lift digging in the lesser blind mole-rat, *Spalax leucodon*; chisel-tooth and scratch digging in most *Ctenomys* spp.).

It is worth noting that there remains uncertainty about digging mode importance in certain lineages. For example, debates over the degree of importance of chisel-tooth digging in some fossorial sciurids have been ongoing for decades (see Burns et al. 1989, Ramos-Lara et al. 2014, Gomes Rodrigues and Damette 2022). To address this issue, we integrated information from multiple sources and assigned a ranking of at least 2 if a given digging mode was consistently described as being typical for a species, particularly if primary behavioral or ecological data were presented or cited. Additionally, “snout-digging” and “head-lift digging” were regarded as equivalent in this study because of the similarities of these movements and the frequent interchangeability of these terms in the literature (Lacey 2000, Hopkins and Davis 2009).

### Quantification of suture complexity

Because suture morphology can be highly complex and variable within and across taxa, more commonly used methods of quantifying shape (e.g., geometric morphometrics, eigenshape analysis) are often not as informative as complexity metrics (White et al. 2020). However, complexity metrics encapsulate shape variation in a single number and vastly differing morphologies may receive similar scores (Allen 2006, White et al. 2020), so it is important to capture multiple axes of shape variation by utilizing several complexity metrics. Therefore, we use three complexity metrics as proxies for suture shape: sinuosity index (SI) (Jaslow 1990, Byron et al. 2018, Nicolay and Vaders 2006, Buezas et al. 2017) and a short-time windowed Fourier transform (STFT) with power spectrum density (PSD) (Allen 2006, White et al. 2020, White et al. 2025) and spectral entropy (Powell and Percival 1979) metrics.

Qualitative ordinal ranking of suture complexity (Byron et al. 2023, Blumer et al. 2024) was not used because of the high degree of variability in overall morphology of the examined sutures and lack of clear qualitative comparison in perceived complexity across taxa. Many sampled sutures also do not exhibit clear fractal-like morphology or delineations between “major” and “minor” lobes or interdigitations, so a suture complexity index approach with lobe counting (see Saunders 1995 for usage of this approach in ammonoids) was not feasible. Although fractal dimension can be a useful complexity metric that may capture unique axes of shape variation compared to other metrics (White et al. 2020), it was not used in this study because our photographs had differing scales and thus resolutions. Additionally, multiple studies that utilize both fractal dimension and sinuosity index have demonstrated nearly identical results across the two methods (see Byron et al. 2004, Buezas et al. 2017), and especially at higher complexities, the two metrics likely capture similar axes of shape variation (Oloriz et al. 1997).

SI is defined as arc length (i.e., length of the suture tracing) divided by the linear distance between the endpoints (i.e., chord length). This is the most commonly used and intuitive method of quantifying suture complexity (see Jaslow 1989; 1990, Saunders 1995, Byron et al. 2004, White et al. 2020, Kammerer 2021, Khurelbaatar et al. 2025). PSD was chosen because it tends to capture a different axis of shape variation than SI (White et al. 2020) and include information about where in the suture complexity is concentrated (Allen 2006). Spectral entropy is commonly used in data science for detecting irregularity in time series data, particularly electroencephalograms (EEGs) (Powell and Percival 1979, Inouye et al. 1991, Nunes et al. 2004, Tenev et al. 2025) and therefore was chosen as a proxy for irregularity in suture structure. Other forms of entropy (e.g., approximate entropy, sample entropy) have been used to quantify complexity and irregularity in other fields (see Pincus 2006, 2008, Humeau-Huertier 2018), and such methods should be assessed as proxies for suture complexity and irregularity in future work.

Prior to the STFT, we slid the semilandmarks to minimize bending energy and placed them in generalized Procrustes superimposition using the gpagen function in the geomorph package (Adams and Otarola-Castillo 2013). The Procrustes-superimposed coordinates were then used to run the STFT (White et al. 2020, White et al. 2025). Although sliding sometimes resulted in “shortening” of interdigitation amplitude, especially in sutures with high interdigitation amplitude, and/or introduction of artificial rugosity to smooth edges (Supplementary Fig. 1), Spearman’s rank correlations between PSD and entropy values derived from the slid and non-slid datasets were very close to 1 (Supplementary Fig. 2), so the slid dataset was used in subsequent analyses. Five specimens (two of the lowland paca, *Cuniculus paca*; and three of the capybara, *Hydrochoerus hydrochaeris*) had extremely short premaxillofrontal sutures (less than 1mm in total length) that were not conducive to the STFT, so these sutures were removed from the PSD and spectral entropy datasets.

Spearman’s rank correlations and associated p-values were calculated for complete pairwise observations of skull lengths and suture complexity metrics, as well as for digging mode importance rankings, using the corrplot package (Wei and Simko 2010). Species-level intraclass correlation coefficients (ICC) were calculated for each complexity metric by extracting variance components from linear mixed models in the lme4 package (Bates et al. 2015) and dividing the variance explainable by species identity by the total variance (Muller and Buttner 1994). An ICC value close to 1 indicates low intraspecific variance, whereas a value less than 0.5 indicates greater distances between specimens of the same species than distances between species means.

### Bayesian multilevel modeling

Because digging mode importance rankings are ordered categorical data, rather than discrete categories or continuous data, we used cumulative ordinal Bayesian multilevel regressions built in brms (Burkner 2017, Burkner and Vuorre 2019, Wisniewski et al. 2023) to test our hypotheses. We used importance rankings for each digging mode as our ordinal response variables and the suture complexity metrics as our predictor variables. To account for and examine the potential effect of body size on digging-related function and allometric relationships between body size and suture complexity, we also modeled the interaction of log-transformed skull length and the complexity metrics.

Because intraspecific variance in suture complexity was often high, especially in the nasofrontal suture (Table 1), we used a measurement error term for our predictor variables instead of simple species means. This approach was chosen because all conspecifics had the same response value, so a standard species-level grouping effect would absorb all residual variance and mute real functional signal. A within-between decomposition (see Guo et al. 2021) would also have been unsatisfactory because of potentially skewed estimates of species means as a result of imperfect sampling, whereas a measurement error term can explicitly account for the effect of sample size on uncertainty of species means estimates.

**Table 1:** Species-level intraclass correlation coefficient (ICC) for each metric. ICC values closer to 1 indicate reduced within-species variance, and ICC values closer to 0 indicate higher within-species variance.

| Suture | Metric | ICC Value |
| --- | --- | --- |
| PMF | SI | 0.728 |
| PMF | PSD | 0.578 |
| PMF | Spectral entropy | 0.598 |
| NF | SI | 0.543 |
| NF | PSD | 0.472 |
| NF | Spectral entropy | 0.34 |

Because 11 specimens were missing skull lengths due to damaged occipital regions, these values were imputed using the phylopars function in the Rphylopars package (Goolsby et al. 2015). All continuous variables (i.e., suture complexity metrics and imputed and log-transformed skull lengths) were transformed into z-scores. Species means, variances, and pooled standard deviations for each complexity metric were then obtained, and species-level measurement error was calculated by dividing the pooled standard deviation by the squared sample size per species. Because skull length was not nearly as variable intraspecifically as the complexity metrics and multiple measurement error terms would vastly inflate the complexity of our models, species means for skull length were used. We built our models using a cumulative probit family and accounted for shared evolutionary history between taxa by incorporating a random effect derived from the phylogenetic correlation matrix.

Our models took the general form:

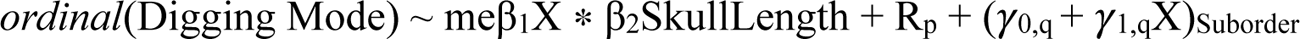

where β1 is the effect of complexity metric X on digging mode importance, β2 is the effect of skull length, β1*β2 is the interaction between X and skull length, and Rp is the phylogenetic random effect derived from the phylogenetic correlation matrix. To allow for explicit comparison of relationships between suture complexity and digging mode importance across suborders, slopes and intercepts were allowed to vary by suborder.

Intercept priors for each digging mode were chosen based on observed rank prevalence. Eight priors from the normal and Student families, as well as an induced Dirichlet prior (see Wisniewski et al. 2023), were tested using prior predictive checks, and the prior that most closely resembled the observed data was adjusted. Priors tested include: normal(0, 1), normal(0, 1.5), normal(0, 2), normal(0, 0.5), student(3, 0, 1), student(3, 0, 2), student(3, −0.5, 1.5), student(3, 0, 3), and the induced Dirichlet prior. Chosen priors include: normal(−0.75, 1) for scratch digging, normal(0.5, 0.75) for chisel-tooth digging, and normal(2.5, 1.5) for head-lift (see Supplementary Fig. 3 for intercept-only prior predictive checks). Normal(0, 1) priors were selected for z-scaled predictors and skull lengths, as well as the phylogenetic correlation matrix. Weakly informative Lewandoski-Kurowica-Joe priors (LKJ = 2) were selected for the correlation of random slopes and intercepts within each suborder (Burkner 2017); although a flat prior (LKJ = 1) and a stronger prior (LKJ = 5) did reshape the posterior distribution of these correlations, suborder-level results were very similar across models with different LKJ priors (see supplementary code). For each model, four chains with 4,000 iterations (2,000 iterations of burn-in and 2,000 of sampling) were run.

### Assessment of model performance and effect sizes

Model convergence and performance were assessed in multiple ways. General model performance and fit were evaluated using posterior predictive check plots in which 500 random draws from the posterior distribution were compared to the empirical counts of the rankings for each response (Supplementary Figs. 4-6). The Gelman-Rubin Rhat statistic was used to confirm adequate chain convergence (Gelman and Rubin 1992). The significance of predictor effects (i.e., the magnitude of the relationship between complexity metrics and predicted digging mode rankings) was assessed using the 89% credible intervals (CIs) of conditional effect sizes. If the 89% CI excluded zero, the effect was considered significant, and if the 89% CI included zero, the effect was considered not significant. Significance of interaction of predictor effects and skull size was estimated similarly: if the 89% CI of the interaction term excluded zero, the interaction was considered significant. Effect size significance for each complexity metric per digging mode was estimated across the entire dataset and within each suborder (Anomaluromorpha, Castorimorpha, Hystricomorpha, Myomorpha, and Sciuromorpha).

For each model that produced a significant predictor effect in at least one suborder, we first evaluated individual model accuracy using the original dataset and the predict function to estimate the posterior probability of a species being assigned to a given rank (Burkner 2017). We then subtracted the predicted rank with the highest posterior probability from the empirical rank and calculated exact accuracy (i.e., predicted rank = empirical rank) and within-1 accuracy (i.e., predicted rank = empirical rank ± 1) for each model. We then used a model stacking approach, as per Wisniewski et al. (2023), to assess the relative performance of each model with significant effect sizes and assign model weights. Because the cumulative ordinal probit structure and measurement error terms produced high per-observation leverage that tended to inflate Pareto-k scores, which are one way of assessing model performance and the influence of individual observations on the posterior distribution (Vehtari et al. 2016), model accuracy was assessed using 10-fold cross-validation instead. A stratified split using the kfold_split_stratified function in the loo package (Vehtari et al. 2015) ensured that each fold contained species with each ranking and that species identities within each fold were equivalent across models with the same response variable. The loo_compare function was then used to evaluate expected predictive accuracy across models via differences in expected log pointwise density (ELPD) before calculating stacking weights (Yao et al. 2018). If a given model performed significantly worse than the others (ELPD difference / standard error > 2) and received a stacking weight of 0, it was excluded from subsequent predictive analyses. If the expected predictive performance of all remaining models was statistically indistinguishable (ELPD difference / standard error < 2), the models were weighted equally to avoid assigning stacking weights based on noise rather than actual differences in accuracy. We calculated model-averaged predicted rank probabilities from 1,000 posterior draws and computed a weighted importance score (WIS) for each draw, rounded each WIS to the nearest integer, and subtracted the rounded WIS from the empirical rank.

### Prediction of digging mode importance in cryptic species

Of the 107 species for which we collected morphological data, 11 had insufficient literature data for one or more digging mode ranking assignments (Fig. 1). Seven species (the mountain degu, *Octodontomys gliroides*; the montane guinea pig, *Cavia tschudii*; the California vole, *Microtus californicus*; the rock pocket mouse, *Chaetodipus intermedius*; the Michoacan pocket gopher, *Zygogeomys trichopus*; the hispid pocket gopher, *Orthogeomys hispidus*; and the giant pocket gopher, *Orthogeomys grandis*) lacked data for all three digging modes, two (the Sonoma tree vole, *Arborimus pomo*; and the red tree vole, *Arborimus longicaudus*) lacked information for scratch digging only, two (the short-tailed bandicoot rat, *Nesokia indica*; and the lesser bandicoot rat, *Bandicota bengalensis*) lacked data for scratch and head-lift digging, and one (the mountain beaver, *Aplodontia rufa*) lacked data for chisel-tooth and head-lift digging. We used our fitted models to predict digging mode importance in these species. To reduce noise, only models with significant effect size(s) and nonzero stacking weights were used for predictions. To test how strongly phylogenetic context informs predictions, we compared phylogenetically naive and phylogenetically conditioned predictions. To obtain phylogenetically naive predictions, we sampled 4,000 posterior draws of the predicted digging mode importance rank value from the posterior distribution of each applicable model and computed a weighted importance score (WIS) for each draw, as per Wisniewski et al. (2023). We then used a likelihood-adjustment approach to obtain phylogenetically conditioned posterior probability estimates. We assigned a placeholder value for the unknown digging mode rankings and performed standard cumulative runs for all 107 species, then subtracted the log probability mass function for the species with missing response data to nullify their net contribution to the model. We subsequently extracted 4,000 predictions from each model, dropping species-level intercepts and measurement error but retaining the phylogenetic term to obtain predictions conditioned on shared evolutionary history, and computed WIS. In cases of uncertain prediction, WIS approaches may result in a regression to the mean effect, but Type I error rates are reduced when compared to maximum *a posteriori* or posterior mode estimates (Wisniewski et al. 2023).

**Fig. 1:**
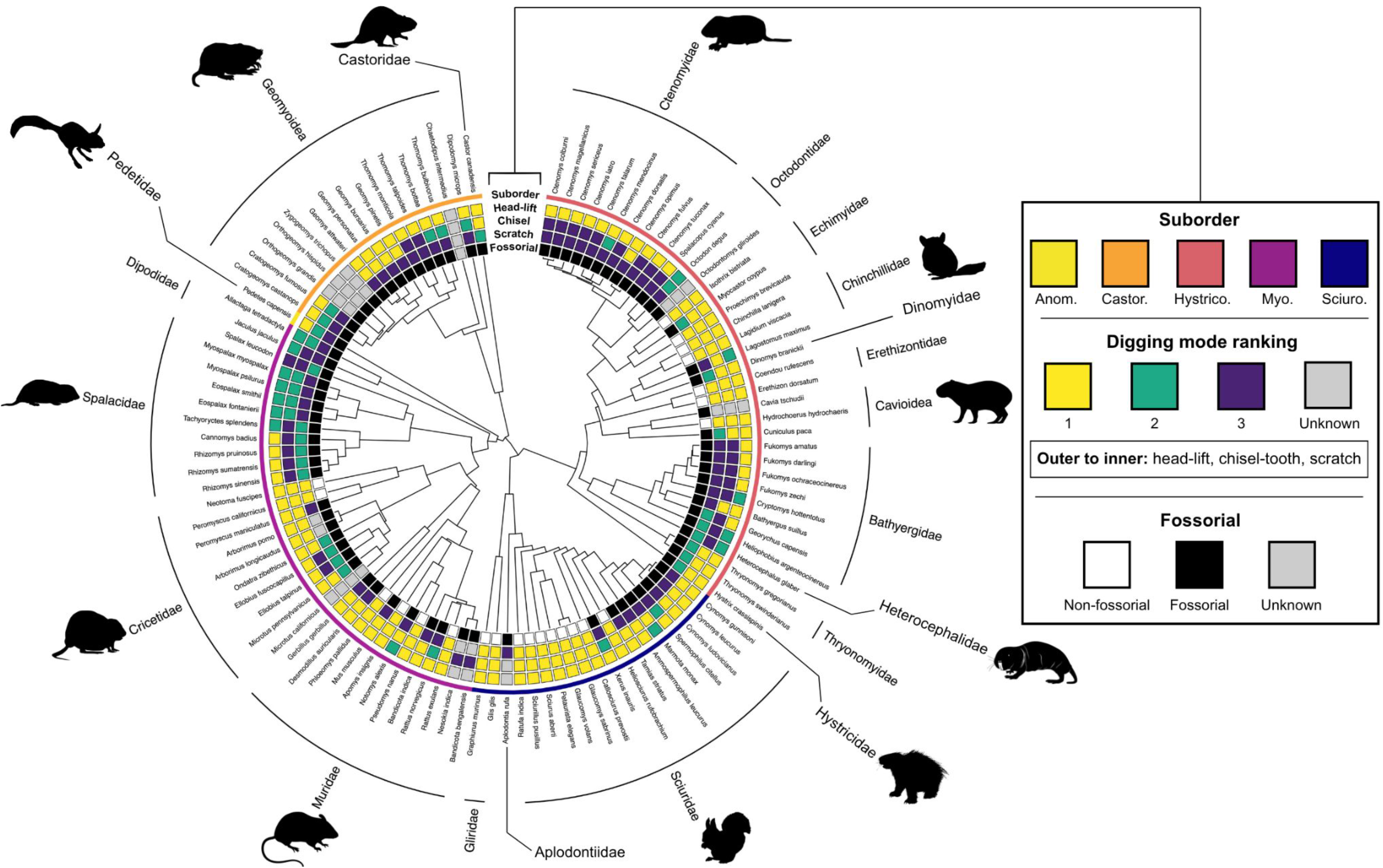
Phylogenetic distribution of digging mode importance rankings for the 107 sampled species, including 11 species that are data-deficient for at least one digging mode. From innermost to outermost, the rings represent binary fossoriality (i.e., whether a taxon regularly digs), scratch digging importance, chisel-tooth digging importance, head-lift digging importance, and suborder. Families are labeled around the circle, and silhouettes are sourced from Phylopic. *Ctenomys minutus* silhouette by Chloe Schmidt; all other silhouettes are openly available under a public domain license.

## Results

### Distribution of digging mode importance rankings

All species known to be fossorial and data-rich for scratch digging (a total of 96 spp.) scored at least a 2 for this digging mode; 23 non-fossorial species were assigned a rank of 1, 20 fossorial species were assigned a rank of 2, and 53 fossorial species were assigned a rank of 3 (Fig. 1). Therefore, it is highly likely that the vast majority of, if not all, fossorial rodents scratch dig, at least in a marginal capacity. Chisel-tooth digging was rarer in the dataset. Of the 99 species assigned a chisel-tooth importance ranking, 16 species received a ranking of 2, and 30 species scored a 3. Head-lift digging was by far the rarest digging mode (Fig. 1). Among the 97 species with assigned head-lift rankings, 14 species were given a ranking of 2, and only one (*Spalax leucodon*) received a ranking of 3 (Fig. 1). Of the species with assigned rankings for all three digging modes, 23 engage only in scratch digging, and 50 species regularly use at least two digging modes, with 12 utilizing all three. Usage of all three digging modes was most common within Dipodidae, Spalacidae, and Bathyergidae. No species received a ranking of 3 for all digging modes. All digging modes had significant positive correlations with one another (Fig. 3A), but the correlation between scratch and chisel-tooth rankings was the strongest (Spearman’s *ρ* = 0.39, p < 0.001).

### Suture complexity metrics

Particularly in the premaxillofrontal suture, the three complexity metrics capture different axes of shape and complexity variation. SI measures how much the suture “meanders” from the Euclidean distance between endpoints, with sutures exhibiting many high-amplitude interdigitations scoring the highest and linear sutures scoring the lowest (Fig. 2). The ceiling for SI values is substantially higher for the premaxillofrontal suture than for the nasofrontal, whereas the ranges of PSD and spectral entropy values are more consistent across the two sutures (Fig. 2). PSD separates sutures with tortuous, looping morphology from those with high-amplitude interdigitations and picks up on general suture orientation (i.e., u-shaped vs. n-shaped) even after Procrustes superimposition because it contains information about where in the curve complexity is concentrated (Fig. 2; see Allen 2006). The effect of suture orientation on PSD values is especially apparent in the nasofrontal suture, which can project more anteriorly or more posteriorly in different species, whereas the general orientation of the premaxillofrontal suture tends to be more consistent across taxa. Spectral entropy separates sutures with a few large, often regularly spaced interdigitations from sutures with more looping, blobular morphology (Fig. 2). It is worth noting that spectral entropy does not always correlate with visual interpretation of irregularity in the raw signal (Anier et al. 2012), but rather is a metric of complexity and irregularity in the frequency domain. The ability of different kinds of entropy (e.g., approximate entropy, joint entropy, sample entropy) to approximate irregularity in the raw suture morphology should be assessed in future work.

**Fig. 2:**
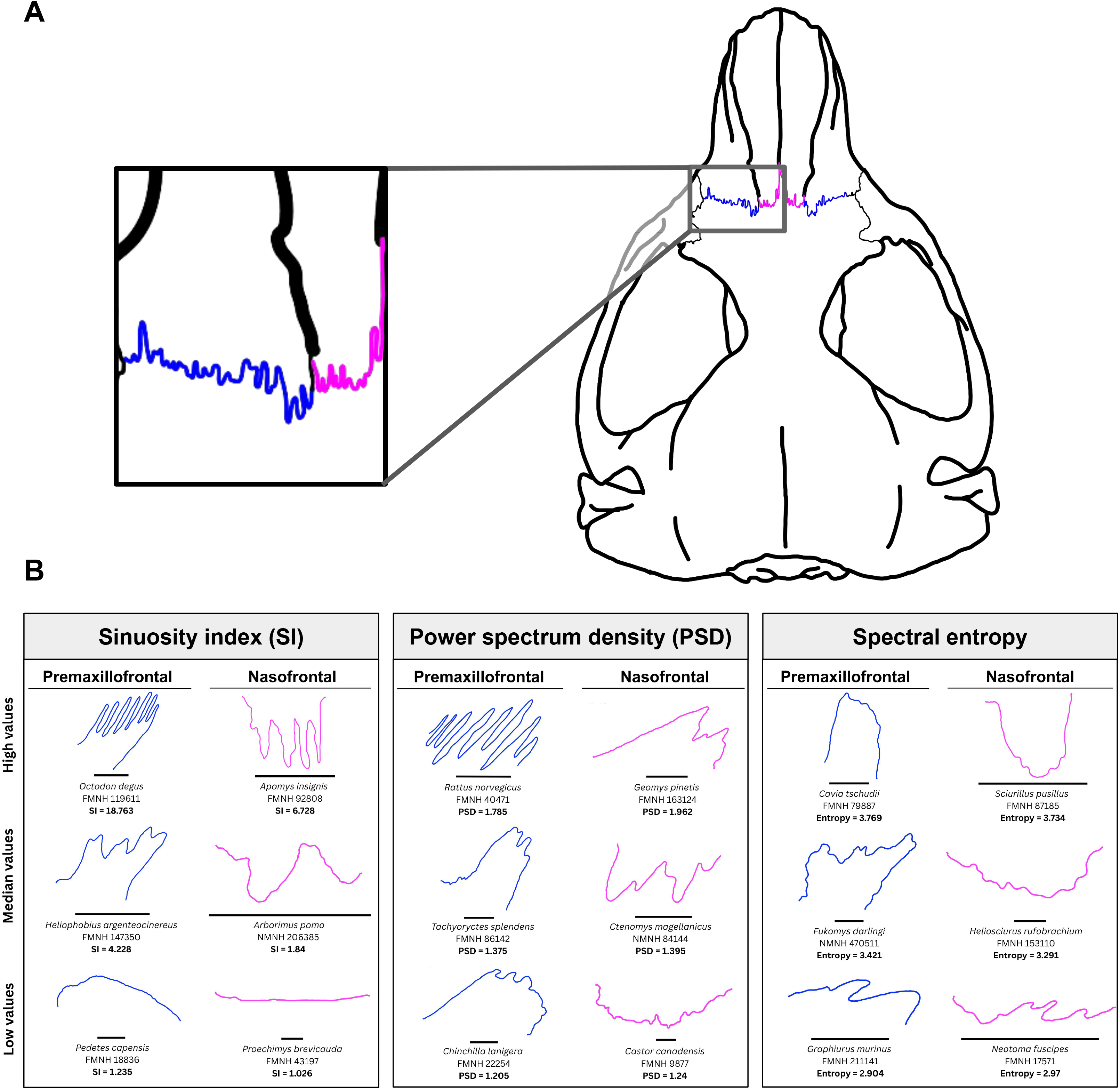
A: Line drawing of the skull of *Aplodontia rufa* (Sciuromorpha: Aplodontiidae) in dorsal view with premaxillofrontal and nasofrontal sutures highlighted and the sampled left side magnified. Specimen ID: FMNH 6320. B: Example premaxillofrontal and nasofrontal sutures that score very high, around the median, and very low for each complexity metric (sinuosity index, power spectrum density, and spectral entropy). Scale bars represent 1mm. Museum abbreviations: Field Museum of Natural History (FMNH), Smithsonian National Museum of Natural History (NMNH). Suture abbreviations: premaxillofrontal (PMF), nasofrontal (NF).

For the premaxillofrontal suture, PSD and spectral entropy are nearly orthogonal (Spearman’s *ρ* = −0.05, p = 0.84), whereas PSD and SI are significantly positively correlated (Spearman’s *ρ* = 0.5, p = 0.012), and spectral entropy and SI are insignificantly positively correlated (Spearman’s *ρ* = 0.22, p = 0.66) (Fig. 3B). The three metrics capture more similar axes of shape and complexity in the nasofrontal suture, with all three being significantly correlated, SI and PSD the most strongly so (Spearman’s *ρ* = 0.7, p = 0.003) (Fig. 3B). Skull length is weakly to moderately negatively correlated with all complexity metrics in both sutures, significantly so with premaxillofrontal SI (Spearman’s *ρ* = −0.4, p = 0.034) and PSD (Spearman’s *ρ* = −0.21, p = 0.048) (Fig. 3B).

**Fig. 3:**
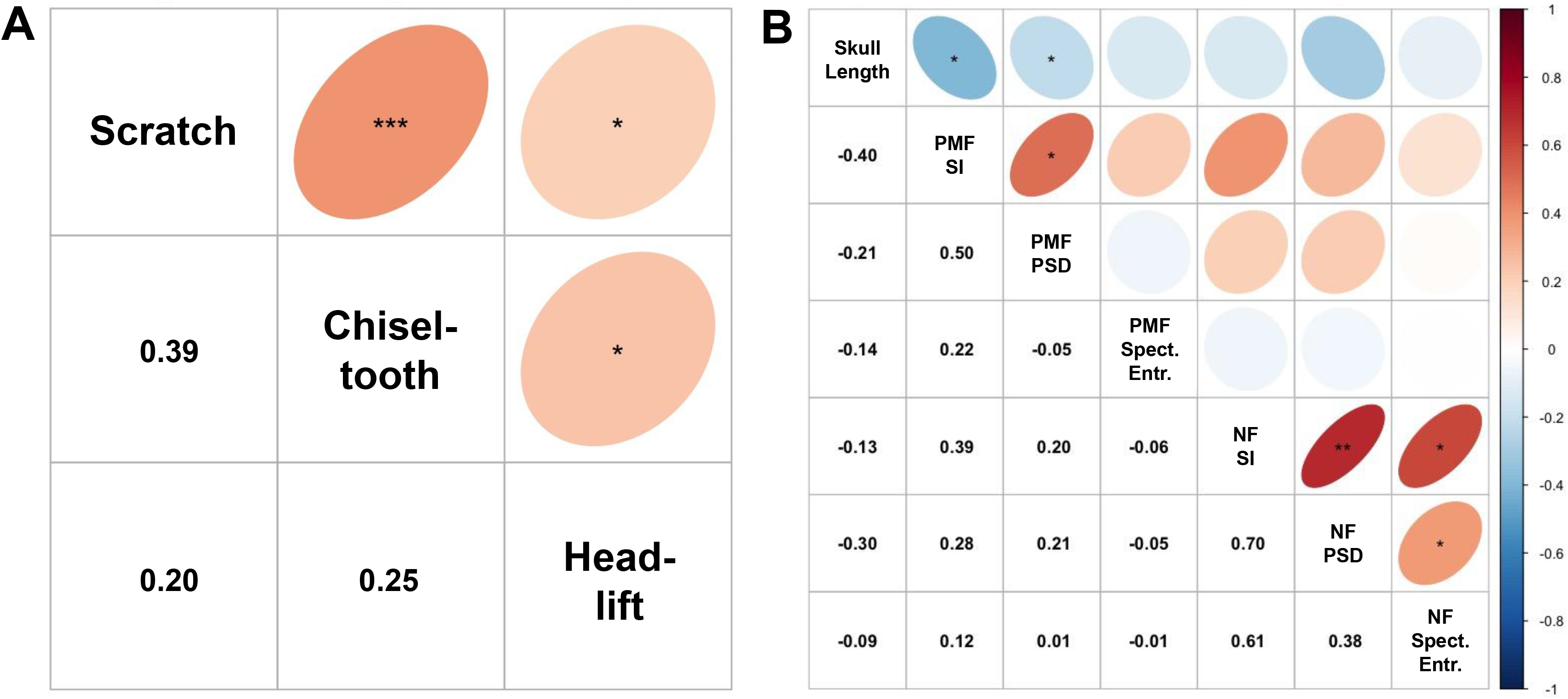
Correlation plots of Spearman’s rank correlations and associated p-values between importance of different digging modes (A) and between complexity metric scores as well as log-transformed skull length (B). P-value abbreviations: * = p < 0.05, ** = p < 0.01, *** = p < 0.001.

The nasofrontal suture consistently has a greater degree of intraspecific variation than the premaxillofrontal suture for each metric (Table 1). Premaxillofrontal SI is the metric with the least amount of within-species variance; nearly three-quarters of the variance in this metric can be explained by species structure (ICC = 0.728), whereas just under half of the total variance in nasofrontal SI can be explained by within-species deviations (ICC = 0.543). Similarly, just over half of the total variance in premaxillofrontal PSD (ICC = 0.578) and spectral entropy (ICC = 0.598) can be attributed to differences between species, whereas the distances between specimens of the same species tend to be greater than distances between species means for nasofrontal PSD (ICC = 0.472) and especially spectral entropy (ICC = 0.34).

Suture complexity, as measured by SI and PSD, is consistently elevated in Muridae. The vast majority of sampled murid species (92.3%) have multiple high-amplitude interdigitations in their premaxillofrontal and/or nasofrontal sutures, a morphology that tends to be rarer in most other families. Although other families can exhibit comparable complexity scores for certain metrics (see premaxillofrontal SI in Octodontidae, nasofrontal PSD in Dipodidae), Muridae stands out for its consistent high complexity (Fig. 4).

**Fig. 4:**
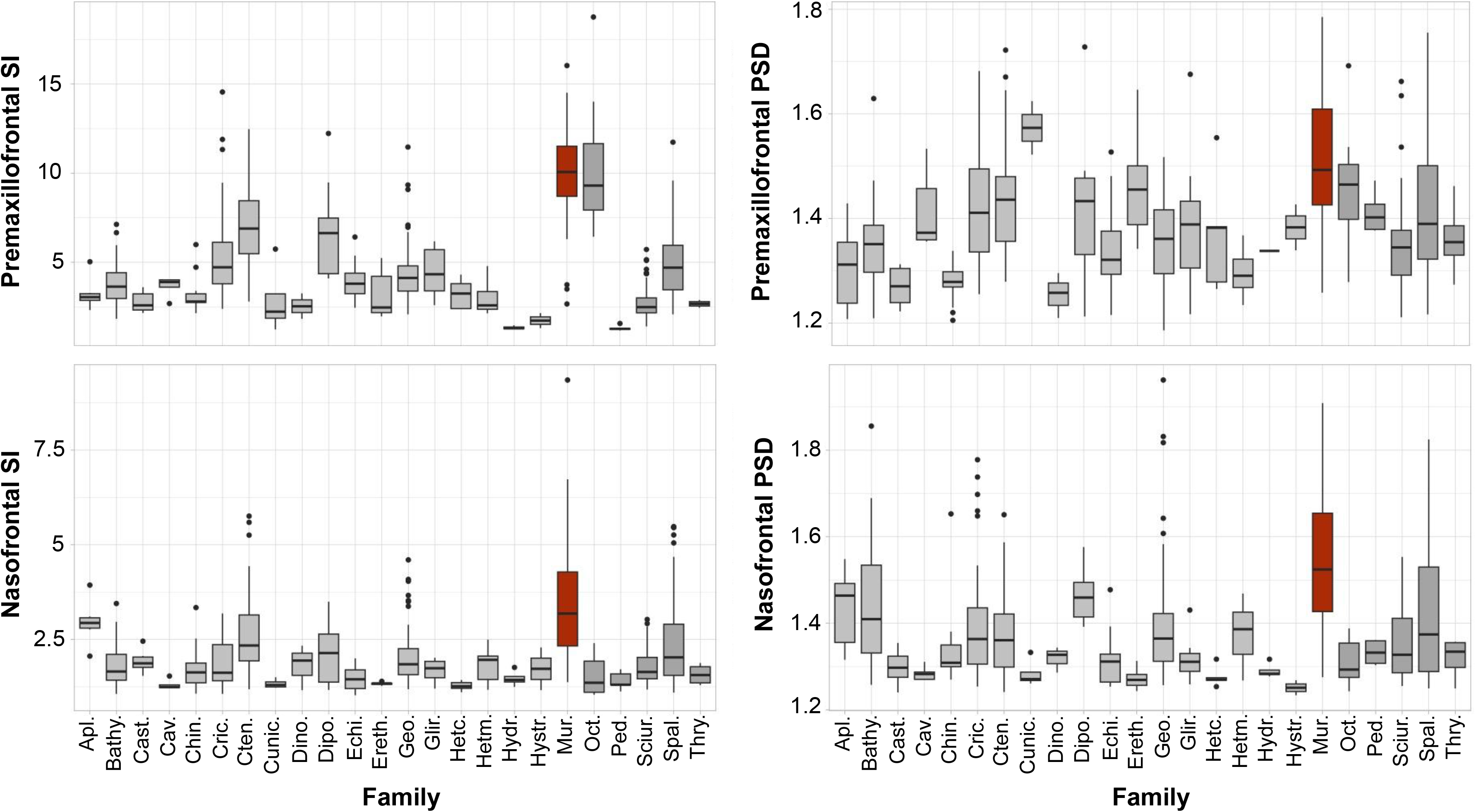
Boxplots of premaxillofrontal SI (upper left), premaxillofrontal PSD (upper right), nasofrontal SI (lower left), and nasofrontal SD (lower right) across each family; Muridae is highlighted in red. Data are at the specimen level. Family abbreviations: Aplodontia (Apl.), Bathyergidae (Bathy.), Castoridae (Cast.), Caviidae (Cav.), Chinchillidae (Chin.), Cricetidae (Cric.), Ctenomyidae (Cten.), Cuniculidae (Cunic.), Dinomyidae (Dino.), Dipodidae (Dipo.), Echimyidae (Echi.), Erethizontidae (Ereth.), Geomyidae (Geo.), Heterocephalidae (Hetc.), Heteromyidae (Hetm.), Hydrochoeridae (Hydr.), Hystricidae (Hystr.), Muridae (Mur.), Octodontidae (Oct.), Pedetidae (Ped.), Sciuridae (Sciur.), Spalacidae (Spal.), Thryonomyidae (Thry.).

### Bayesian multilevel models

Posterior predictive checks for each model closely resembled the observed prevalence of ranks (Supplementary Figs. 4-6), indicating good model fit. Adequate convergence of all model chains was confirmed by consistent Rhat values below 1.002 (Gelman and Rubin 1992). Results differed widely across complexity metrics and digging modes. In the head-lift digging models, 89% credible intervals for all complexity metrics included zero, both across the entire dataset and within each suborder (Fig. 5A). Therefore, it is very difficult to predict head-lift digging importance based on suture complexity metrics.

**Fig. 5:**
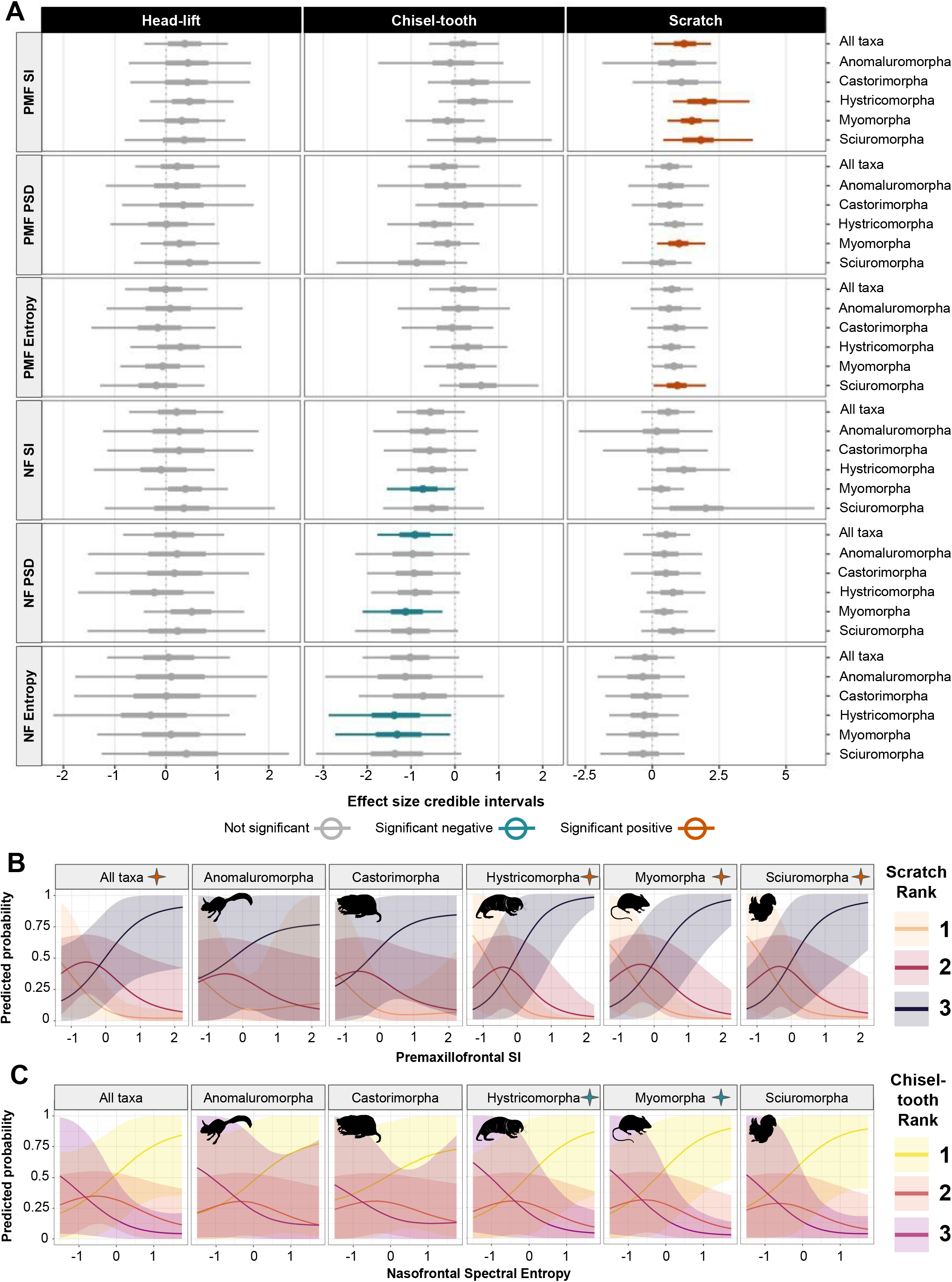
A: 89% credible intervals (CIs) of the posterior conditional effect sizes of each suture complexity metric on digging mode importance rank for the entire dataset (“grand mean”) and each suborder. An effect is considered significant if its 89% CI excludes 0. Suture complexity metrics are listed on the left side of the y-axis, and data subsets are listed on the right side of the y-axis. Teal hues represent models with significant negative effects; orange hues represent models with significant positive effects. Suture abbreviations: premaxillofrontal (PMF), nasofrontal (NF). B: Conditional effects with 89% CIs of premaxillofrontal sinuosity index (SI) on predicted probability of scratch digging importance rank. Y-axis represents predicted probability, x-axis represents z-transformed premaxillofrontal SI value, and panels represent these effects in the entire dataset (far left) and in each suborder (ordered alphabetically). C: Conditional effects with 89% CIs of nasofrontal spectral entropy on predicted probability of chisel-tooth digging importance rank. Y-axis represents predicted probability, x-axis represents z-transformed nasofrontal spectral entropy value, and panels represent these effects in the entire dataset (far left) and in each suborder (ordered alphabetically). All silhouettes are sourced from Phylopic under a public domain license.

Effect sizes tended to be stronger for chisel-tooth and scratch digging. No metric had a significant effect on all suborders, although several metrics had significant effects on multiple suborders and/or the dataset as a whole. Signal in chisel-tooth digging was negative, concentrated in the nasofrontal suture, and strongest in Myomorpha. Nasofrontal SI had a significant negative effect in Myomorpha only, PSD had significant negative effects in the dataset as a whole and in Myomorpha, and spectral entropy had significant negative effects in Myomorpha and Hystricomorpha (Fig. 5A,C). No metrics for the premaxillofrontal suture had significant effects on chisel-tooth rank (Fig. 5A).

All complexity metrics for the premaxillofrontal suture had positive significant effects on scratch digging rank in at least one suborder, indicating that specialized scratch diggers tend to have more complex sutures. Premaxillofrontal SI was the only metric to have a significant effect on scratch digging rank across the dataset as a whole, and it was also significant in Hystricomorpha, Myomorpha, and Sciuromorpha (Fig. 5B). Premaxillofrontal PSD was significant in Myomorpha, and premaxillofrontal spectral entropy had a significant effect in Sciuromorpha (Fig. 5A). Additionally, these relationships between suture complexity and scratch digging tended to persist when binary chisel-tooth and head-lift usage were included as covariates, whereas some of the effects of suture complexity on chisel-tooth rank vanished when accounting for scratch and chisel-tooth usage (Supplementary Fig. 7). Notably, no metric had significant effects in the same suborder across different digging modes.

In all models with significant effect sizes for at least one suborder, exact accuracy ranged from 82.8% to 99% of species predicted correctly, and within-one accuracy was consistently at least 99% (i.e., very few species with one extreme rank were predicted incorrectly as the other extreme) (Table 2). In the stacked models, accuracy was consistently high; for the stacked scratch digging model, 94.8% of species were predicted correctly, for the stacked chisel-tooth digging model, 90.9% of species were predicted correctly, and for both stacked models, within-one accuracy was 100% (Table 2).

**Table 2:** Exact and within-one-rank accuracy scores for all models that have a significant relationship (i.e., the 89% credible interval of effect size excludes 0) between suture complexity and digging mode importance in at least one suborder, as well as the stacked model for each digging mode, averaged across all species. Accuracy scores were calculated as the difference between the observed rank and a weighted average of the distribution of posterior probabilities for each rank per species.

| Digging mode | Model | Exact Accuracy | Accuracy w/in 1 |
| --- | --- | --- | --- |
| Scratch | PMF SI | 0.99 | 1 |
| Scratch | NF SI | NA | NA |
| Scratch | PMF PSD | 0.979 | 1 |
| Scratch | NF PSD | NA | NA |
| Scratch | PMF spectral entropy | 0.958 | 1 |
| Scratch | NF spectral entropy | NA | NA |
| Scratch | Stacked (PMF SI, PSD, and spectral entropy) | 0.948 | 1 |
| Chisel-tooth | PMF SI | NA | NA |
| Chisel-tooth | NF SI | 0.828 | 0.99 |
| Chisel-tooth | PMF PSD | NA | NA |
| Chisel-tooth | NF PSD | 0.869 | 1 |
| Chisel-tooth | PMF spectral entropy | NA | NA |
| Chisel-tooth | NF spectral entropy | 0.869 | 1 |
| Chisel-tooth | Stacked (NF SI, PSD, and spectral entropy) | 0.909 | 1 |

The relationships between suture complexity and digging mode importance were rarely mediated by body size, and never so with scratch digging as the response. Premaxillofrontal PSD had a significant positive interaction with skull length when chisel-tooth rank was the response, and premaxillofrontal spectral entropy significantly negatively interacted with skull length with head-lift rank as the response. To visualize these interactive effects, we followed Wisniewski et al. (2023) and plotted the conditional posterior probabilities of belonging to each rank for values of these traits at very low (−2 SD), low (−1 SD), mean, high (+1 SD), and very high (+2 SD) skull sizes. At below-average skull sizes, increased usage of chisel-tooth digging was associated with reduced premaxillofrontal PSD, but at above-average skull sizes, particularly in very large taxa, this pattern reversed somewhat (Fig. 6A). At very small skull sizes, elevated premaxillofrontal spectral entropy values correlated with increased specialization for head-lift digging, whereas low head-lift importance was predicted for most other body size bins regardless of morphology (Fig. 6B).

**Fig. 6:**
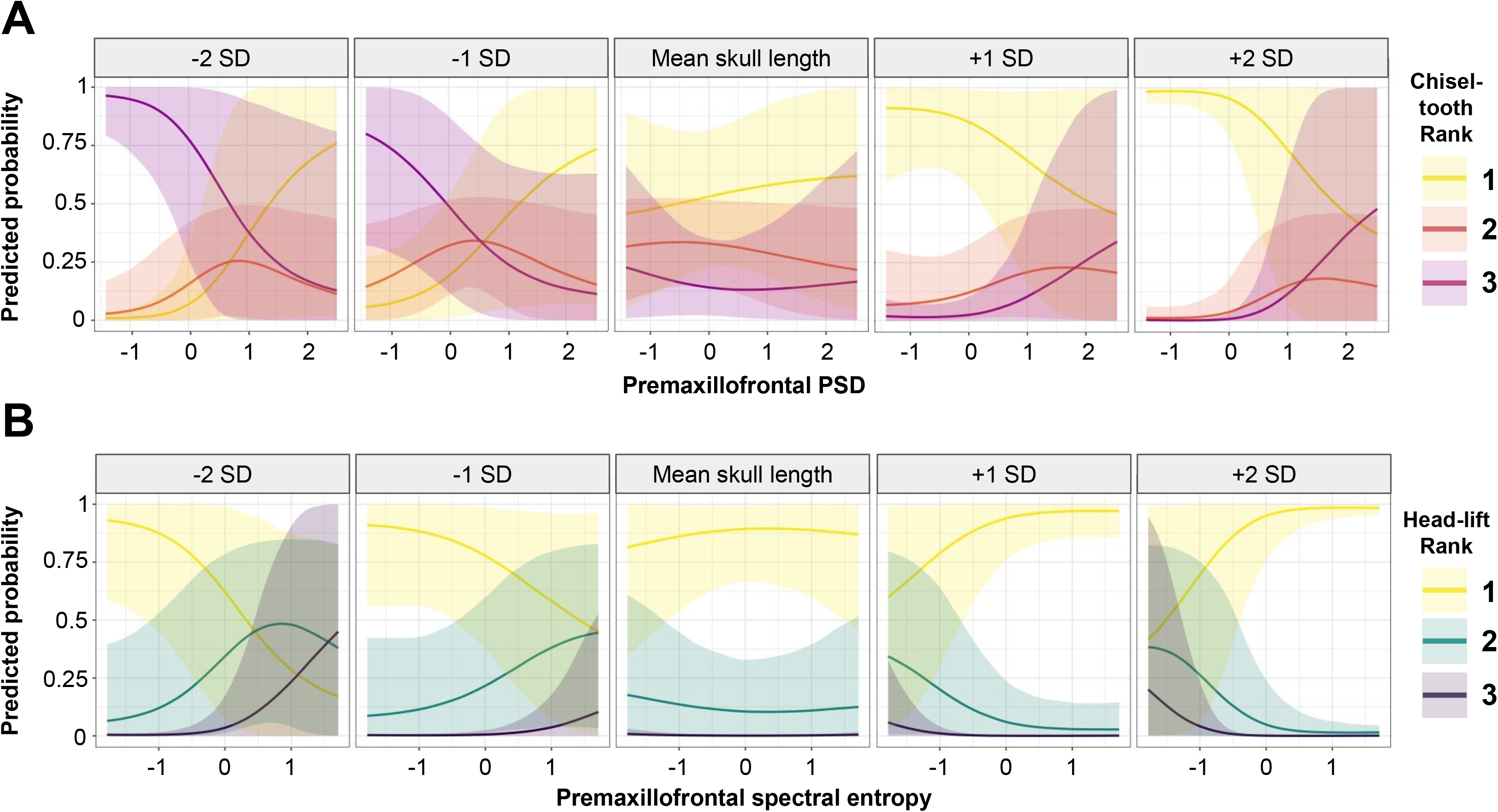
Conditional effect plots demonstrating how skull length mediates the relationship between premaxillofrontal PSD and predicted chisel-tooth importance rank (A) and between premaxillofrontal spectral entropy and predicted head-lift importance rank (B). For each metric, results are shown for species ranging from very small skull length to very large skull length (−2, −1, 0, +1, and +2 standard deviations from the mean, from left to right).

### Prediction of digging mode importance in cryptic species

Because no metrics had significant effects on head-lift digging, we did not predict head-lift rank and instead focused on predictions of scratch and chisel-tooth importance in data-poor species. Predictive accuracy of scratch and chisel-tooth models with significant effect(s) in at least one suborder was high (see above).

When scratch digging rank was the response variable, the performance of the three models with significant effect sizes (premaxillofrontal SI, PSD, and spectral entropy) was statistically indistinguishable (Table 3), so these models were assigned equal weights for subsequent stacking and predictions. The three chisel-tooth models with significant effects (nasofrontal SI, PSD, and spectral entropy) performed approximately equally well (Table 3), so they were similarly assigned equal weights.

**Table 3:** Expected log pointwise density (ELPD) differences and associated standard errors (SE) between models with significant relationships between suture complexity and digging mode importance in at least one suborder, per digging mode. A model is considered to perform significantly worse than the best-performing model if the value of its ELPD difference divided by SE > 2. The performance of models whose ELPD differences / SE < 2 are considered to be statistically indistinguishable.

| Digging Mode | Model | ELPD difference | SE difference |
| --- | --- | --- | --- |
| Scratch | PMF SI | 0 | 0 |
| Scratch | PMF PSD | -5.1 | 2.7 |
| Scratch | PMF spectral entropy | -2.1 | 3.7 |
| Chisel-tooth | NF SI | -1.4 | 2.6 |
| Chisel-tooth | NF PSD | -0.3 | 3.2 |
| Chisel-tooth | NF spectral entropy | 0 | 0 |

Degree of similarity between phylogenetically naive and phylogenetically conditioned digging mode ranking predictions differed across species. Phylogenetically naive predictions of scratch digging importance were relatively homogeneous, with mean WIS ranging from 2.09 (*Chaetodipus intermedius*) to 2.61 (*Nesokia indica*) (Fig. 7). Despite the higher predictive accuracy of the models with scratch digging as the response (see above), this homogeneity suggests greater predictive uncertainty and possibly a regression to the mean effect. The very fossorial *Bandicota bengalensis* (mean WIS = 2.43) and *Nesokia indica*, both of which are murines highly specialized for chisel-tooth digging, received the highest mean WIS for scratch digging out of the eleven species data-deficient for this digging mode. Phylogenetically conditioned predictions for scratch digging were more heterogeneous, with *Microtus californicus* (mean WIS = 1.96) and *Cavia tschudii* (mean WIS = 1.94) having the lowest predicted rankings and the castorimorphs having the highest mean WIS (2.9 for *Zygogeomys trichopus* and *Orthogeomys hispidus*, 2.86 for *Orthogeomys grandis*, and 2.66 for *Chaetodipus intermedius*). The four castorimorph species, as well as *Octodontomys gliroides*, had much higher mean WIS for the phylogenetically conditioned predictions, whereas phylogenetic conditioning slightly decreased mean WIS for *Cavia tschudii* and *Microtus californicus*. The remaining myomorph species had comparable mean WIS across both prediction methods.

**Fig. 7:**
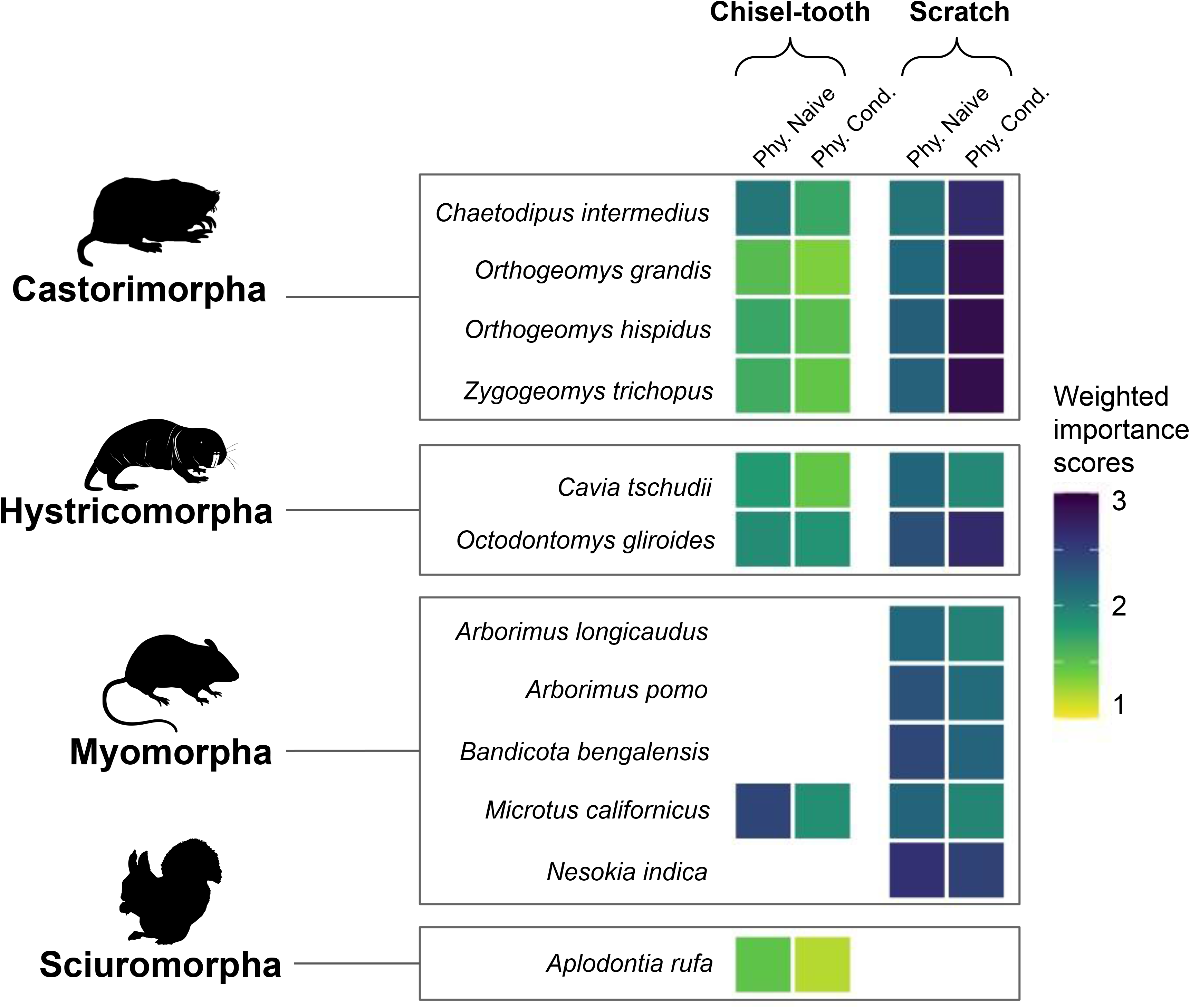
Heatmap of posterior mean weighted importance score (WIS) estimates for chisel-tooth and scratch digging for eleven species data-deficient for at least one of these digging modes. Phylogenetically naive predictions account for morphology only, whereas phylogenetically conditioned predictions account for shared evolutionary history with data-rich species. Color represents mean WIS value, with darker colors corresponding to higher predicted importance scores. Only models with significant effect sizes in at least one suborder were used for these predictions, and they were weighted by comparing expected log pointwise predictive density (see Methods).

Chisel-tooth digging importance predictions were more heterogeneous, especially when the phylogenetically naive method was used, suggesting less predictive uncertainty despite the slightly lower accuracy of these models (see above) (Fig. 7). Mean WIS scores for chisel-tooth digging tended to be lower overall than for scratch digging (Fig. 7). *Aplodontia rufa* received the lowest mean WIS for chisel-tooth digging importance (1.47 using the phylogenetically naive method and 1.19 using the phylogenetically conditioned method), followed closely by the three geomyid species: *Orthogeomys grandis* (mean WIS = 1.52 naive of phylogeny, 1.34 accounting for phylogeny), *Zygogeomys trichopus* (mean WIS = 1.66 naive of phylogeny, 1.45 accounting for phylogeny), and *Orthogeomys hispidus* (mean WIS = 1.7 naive of phylogeny, 1.51 accounting for phylogeny). Only the heteromyid *Chaetodipus intermedius* (mean WIS = 2.08) and the arvicoline cricetid *Microtus californicus* (mean WIS = 2.43) received average predicted chisel-tooth importance of greater than 2 for the phylogenetically naive predictions, but no species received a mean WIS greater than 2 for the likelihood-adjusted predictions. *Microtus californicus* (mean WIS = 1.89) and *Octodontomys gliroides* (mean WIS = 1.85) received the highest predicted rankings, followed closely by *Chaetodipus intermedius* (mean WIS = 1.7) using the latter method (Fig. 7). Predictions for the geomyid species and *Octodontomys gliroides* were similar across both methods, but *Chaetodipus intermedius*, *Cavia tschudii*, *Aplodontia rufa*, and *Microtus californicus* had substantial differences (Fig. 7).

## Discussion

### Drivers of suture complexity

Excavation of earthbound substrate and usage of the subterranean habitat are common aspects of function and ecology across Mammalia and are especially prevalent within Rodentia. More specialized fossorial rodents exhibit remarkable convergence in overall lifestyle and general morphology, including fusiform bodies, short and stout limbs, and reduced eyes and pinnae (see Nevo 1979, Lacey 2000). However, lineages can differ widely in digging-related function, including usage and relative importance of different skeletal elements and movements (i.e., digging modes) for excavation. Rodent species can use up to three digging modes that involve various aspects of the craniodental apparatus and postcranial skeleton and can exhibit varying degrees of specialization for each (Fig. 1). Although other mammalian groups tend to be more constrained in terms of digging mode usage (e.g., all fossorial carnivorans scratch dig, all extant talpids use humeral-rotation), their members also exhibit substantial variation in degree of specialization for digging. For instance, within Carnivora, both red foxes (*Vulpes vulpes*) and American badgers (*Taxidea taxus*) scratch dig, but the latter is clearly far more specialized for both a generally fossorial lifestyle and for this particular digging mode. Fossorial function is a complex, multivariate trait and should be treated as such in comparative studies.

Therefore, we utilized an ordinal ranking system for digging mode importance and built Bayesian hierarchical models that account for within-species variation and shared evolutionary history to examine the relationship between fossorial function and cranial suture complexity in rodents. Suture complexity correlates with aspects of rodent fossoriality, specifically with importance of scratch and chisel-tooth digging but not head-lift digging. However, no complexity metric predicts importance of multiple digging modes across all rodent suborders. Instead, these relationships vary substantially across digging modes and clades, suggesting that both phylogenetic constraints and usage of disparate elements and movements during substrate excavation place different selective pressures on specific axes of suture morphology. These suborder-specific trends in part may be due to shared developmental constraints or ecology-independent selective pressures that affect suture morphology more strongly than species- or genus-level fossoriality-related function and ecology.

Surprisingly, the strongest and most consistent relationship between suture morphology and fossorial function is between premaxillofrontal suture complexity and scratch digging importance. Each premaxillofrontal complexity metric had significantly positive effects in at least one suborder, most commonly in Myomorpha (SI and PSD) and Sciuromorpha (SI and spectral entropy) (Fig. 4). This positive signal may be partially explained by more specialized fossorial species’ harder diets, which often consist of fibrous geophytes and/or phytolith-rich grasses (Fagerstone et al. 1981, McIntosh and Cox 2016a, Augustine et al. 2023). Specialized scratch diggers are also more likely to engage in other digging modes, at least secondarily (Fig. 1, Fig. 2A). However, when chisel-tooth and head-lift digging usage were included as binary covariates, the signal remained (Supplementary Fig. 7), suggesting that this phenomenon is unique to scratch digging, rather than due to dietary differences or implicit correlation with other digging modes. It is possible that during scratch digging, especially in more specialized taxa that tend to be smaller-bodied and shorter-limbed, the snout may be pressed against the substrate for balance or leverage. Such behavior would likely expose the craniofacial region to compressive strains, which could explain the more complex suture morphology observed in more specialized scratch diggers. Description of this type of behavior is lacking in the literature, and examination of head and snout kinematics during scratch digging, especially across distantly related species, is needed to test this hypothesis.

Also contrary to our H1, nasofrontal—but not premaxillofrontal—suture complexity tended to be negatively correlated with chisel-tooth digging importance, especially in Myomorpha. SI had a significant effect only in Myomorpha. Given the deeply interdigitated suture morphology observed in chisel-tooth digging caviomorphs (Octodontidae and Ctenomyidae; see Buezas et al. 2017), it is surprising that this effect is not significant in Hystricomorpha as well. However, chisel-tooth digging phiomorphs (most non-*Bathyergus* bathyergids and *Heterocephalus glaber*) frequently have less deeply interdigitated sutures than octodontids and ctenomyids. Their premaxillofrontal sutures tend to have more scalloped or “branching” structure rather than multiple high-amplitude interdigitations (see Fig. 2B for examples), and their nasofrontal sutures tend to be quite simple, with few interdigitations. This stark difference in morphological structure across hystricomorph groups with similar fossorial function may help explain the lack of signal in SI within this suborder, at least with regard to chisel-tooth digging importance.

The negative effect of PSD on chisel-tooth rank in Myomorpha, which drives its significance in the dataset as a whole, likely reflects more posteriorly projecting nasofrontal suture morphology (i.e., u-shaped rather than n-shaped when the specimen is viewed dorsally and facing upward, as per Fig. 2) among more specialized chisel-tooth diggers. Myomorpha and Sciuromorpha also exhibit reduced spectral entropy associated with chisel-tooth specialization, indicating that multiple distinct frequency spikes, rather than more tortuous, blobular morphology, have adaptive significance for this digging mode. Chisel-tooth digging may expose the nasofrontal suture to more tensile strains, particularly in Myomorpha, thus resulting in overall simpler morphology. However, load paths and strain patterns during chisel-tooth digging are not well understood, especially in a comparative context. For instance, Van Wassenbergh et al. (2017) described the kinematics of chisel-tooth digging in *Fukomys micklemi* but acknowledged that other species, even within Bathyergidae, likely exhibit different kinematic patterns than *F. micklemi*. Kinematic variation at the species level and above may result in different taxa experiencing different types of biomechanical strains, which could explain phenomena such as the apparent lack of SI signal in hystricomorphs and the marked disparity in craniofacial suture morphology between chisel-tooth digging caviomorphs and phiomorphs. Finite element analysis focused on the effect of gape on skull performance in bathyergids revealed that greater gape increases biting efficiency but also shifts forces toward the occiput (McIntosh and Cox 2016b), which might also help explain why some chisel-tooth digging groups have less complex rostral sutures. Nevertheless, the general and species-level impacts of suture morphology on skull performance during chisel-tooth digging remain enigmatic.

The lack of signal in head-lift digging in part may be due to the relative rarity of this digging mode, as only 15.46% of data-rich species received a head-lift rank of 2 or above, and only *Spalax leucodon* received a rank of 3. Especially outside of Spalacidae and Dipodidae, usage of head-lift digging is sparse and not particularly phylogenetically conserved, a pattern which may substantially challenge our models. Very little is known about the load paths, kinematics, and strain regimes associated with head-lift digging in rodent lineages, but the non-significant relationship between head-lift importance and suture complexity might also be explained by the presence of both tensile and compressive forces during excavation. Nicolay and Vaders (2006) found that male white-tailed deer (*Odocoileus virginiana*) do not exhibit elevated suture complexity despite their large antlers and interlocking behavior, possibly due to the fact that antler interlocking can generate both tensile and compressive forces, creating a “confusing” environment for suture growth. As with chisel-tooth digging, kinematic and biomechanical studies are needed to test this hypothesis in head-lift diggers. It is also worth noting that although head-lift digging trogonophid amphisbaenians exhibit increased complexity in their transverse craniofacial sutures, the similarly digging Florida worm snake (*Rhineura floridiana*) does not have such complex sutures (Gans 1960), suggesting a clade-specific, nuanced relationship between head-lift digging and suture complexity in a non-mammalian study system. *Rhineura floridiana* tends to inhabit softer, sandier soils than trogonophids (Hipsley and Muller 2014), so this difference may reflect the impact of substrate properties on degree and type of biomechanical strain experienced. Differences in soil properties, as well as dietary hardness and other environmental and ecological factors, therefore should be considered in future work on suture complexity and digging mode usage within and outside of Mammalia.

The relationship between fossorial function and suture morphology is generally strongest and most consistent within Myomorpha. Biting efficiency is higher in rats than in squirrels or guinea pigs (Cox et al. 2012), suggesting an inherent functional advantage of the myomorph zygomasseteric configuration over the hystricomorph and sciuromorph configurations. Therefore, it is possible that lineages within Myomorpha generally experience less feeding-related constraint on the masticatory apparatus than other suborders and are able to allocate more of their craniodental morphology, including suture morphology, to other purposes, such as digging. However, this explanation is highly speculative and needs empirical testing. In contrast, Anomaluromorpha and Castorimorpha tend to exhibit very weak correlations between suture complexity and digging mode importance. In the case of the former, this lack of signal is likely due to the presence of only one representative species (*Pedetes capensis*) in the dataset. Castorimorpha has a relatively sparse presence (n = 12 species, four of which completely lack digging mode importance data) compared to other suborders (n = 39 species for Hystricomorpha, n = 38 species for Myomorpha, n = 18 species for Sciuromorpha), which could contribute to its weak signal. The data-rich sampled castorimorph species tend to be more functionally homogeneous than other suborders (e.g., all geomyoid species were assigned a rank of 3 for scratch digging), which could also obscure fossoriality-related adaptive significance of suture complexity in this group.

The consistently elevated suture complexity among Muridae regardless of fossorial behavior (Fig. 4) has no obvious functional or dietary correlate. Most other myomorph species do not exhibit comparable deeply interdigitated morphology or levels of complexity, so this phenomenon is likely not due to biomechanical strains brought about by the myomorph zygomasseteric configuration. Additionally, several species with extreme complexity (e.g., the Polynesian rat, *Rattus exulans*) tend to rely on softer foods such as termites, fleshy fruits, and earthworms (Fall et al. 1971). Instead, homologous biomechanical stresses or developmental constraints may contribute to this phenotype. The rostral region has been noted as the location with the most strongly developed suture interdigitations in lab mice and rats, possibly because it is subjected to the most prolonged growth period and therefore unique exposure to forces of expansive and adjustive growth (Massler and Schour 1951, Moss 1957), but it is unclear whether this pattern extends to most or all murids. Additionally, finite element analysis (FEA) of lab rat skulls reveals that the nasofrontal suture in particular experiences increased compressive strain during incisor biting (Sharp et al. 2023), so murids may experience unique strain during biting. FEA-based studies that explicitly examine the role of cranial sutures outside of lab rats are extremely rare, so it is uncertain whether the nasofrontal and/or premaxillofrontal sutures experience similar strain intensity across or even outside of Muridae. If the latter is the case, it might explain why rodents tend to have higher transverse craniofacial suture complexity than many other mammalian clades, including Carnivora, Eulipotyphla, Afrotheria, Perissodactyla, Xenarthra (Strassberg, personal observation), and most Artiodactyla (see Herring 1972 for a description of comparable nasofrontal complexity among some suoid species), but would not explain why murids exhibit uniquely elevated complexity compared to other rodents.

We found high degrees of intraspecific variation in suture complexity, especially within the nasofrontal suture and the STFT-derived metrics. Individual-level differences in diet, disparity in exposure to biomechanical stresses (digging-related or otherwise), or general lifestyle variation may contribute to this phenomenon. However, it is impossible to perfectly account for individual-level variation in factors such as dietary hardness, amount of time spent using different digging modes, and precise soil qualities, especially in wild-caught animals. Even accession or field notes that convey information regarding stomach contents, trapping location, or behavior between capture and death represent a mere snapshot and may not be indicative of an individual’s overall lifestyle. Studies that explicitly integrate digging-related behavioral and kinematic data, ontogeny, and suture morphology are needed to better understand the mechanisms underlying intraspecific variation in suture complexity, particularly in species that are not commonly used for suture studies. *In vivo* research on biomechanical stresses and suture morphology is relatively common in lab mice, lab rats, some primate species, and miniature pigs (e.g., Moss 1957, Herring 1972, Byron et al. 2004, Byron et al. 2023), but relationships between forces and suture morphology are largely unknown outside of these taxa.

It is worth noting that numerous specimens in the dataset exhibit at least a small degree of asymmetry in aspects of suture morphology such as number, amplitude, and/or shape of interdigitations. The roles of development, evolutionary history, function, and ecology in shaping the presence and magnitude of such asymmetry should be considered in future work.

### Suture complexity as a predictive tool

We used models with significant effect size(s) in at least one suborder to predict scratch and chisel-tooth digging importance in 11 data-poor extant species. To evaluate how strongly morphology and phylogenetic context each contribute to predictions, we used both a phylogenetically naive approach and a likelihood adjustment approach that conditioned predictions on the phylogenetic correlation matrix (see Methods). Mean WIS for chisel-tooth digging tended to be lower than those for scratch digging, possibly reflecting the overall lower prevalence of chisel-tooth digging (Fig. 1, Fig. 7). Phylogenetically naive chisel-tooth ranking predictions were more heterogeneous than the phylogenetically conditioned predictions, whereas the opposite was true for predictions of scratch digging importance, possibly indicating that phylogenetic context increased uncertainty for chisel-tooth but not scratch digging. For some species, predicted digging mode importance was similar across both methods (e.g., scratch digging importance in *Arborimus longicaudus*, *Arborimus pomo*, *Bandicota bengalensis*, and *Nesokia indica*, chisel-tooth digging importance in *Octodontomys gliroides* and the three geomyid species), suggesting that suture morphology has good predictive power in these cases. However, in other cases, most notably scratch digging importance in the data-poor castorimorph species and *Octodontomys gliroides* and chisel-tooth importance in *Microtus californicus*, phylogenetic conditioning substantially changed mean WIS values, indicating high weight given to shared evolutionary history compared to morphology. In these scenarios, phylogenetic conditioning resulted in predictions more similar to the observed digging mode importance rankings in the data-poor species’ closest relatives. The overall weak functional signal and functional homogeneity in data-rich Castorimorpha (see above) in part may explain the dramatic changes in predicted ranking values for data-poor geomyoids. Infrequent usage of chisel-tooth digging outside of Ellobiusini within Cricetidae similarly might explain why phylogenetically conditioned predictions resulted in much lower mean WIS values for *Microtus californicus*. However, because digging mode importance can be apomorphic and generally tends to be evolutionarily labile in rodents (Fig. 1), it is possible that phylogenetic conditioning may obscure meaningful functional signal in certain contexts.

Given the inferred evolution of head-lift and possibly chisel-tooth digging in extinct derived mylagaulid aplodontiids (Hopkins 2005) and the notable lack of information regarding non-scratch digging mode usage in the extant *Aplodontia rufa*, predictions for the latter taxon were of particular interest. Some aspects of this species’ cranial morphology (e.g., triangular skull outline, broadened occipital region), could be interpreted as adaptations for craniodental digging modes (Hopkins 2005), but others (e.g., dorsoventrally flattened skull with low occiput) suggest otherwise. Regardless of whether phylogenetic context was used for predictions, *Aplodontia rufa* received the lowest mean WIS for chisel-tooth digging of the eight species data-deficient for this digging mode (Fig. 7). These results suggest poor ability to utilize the incisors for soil excavation. Little is known about digging mode usage in fossil Aplodontinae; it is possible that chisel-tooth digging ability was secondarily lost or simply never evolved in the lineage leading to *Aplodontia rufa*. Study of behavior and kinematics in live *Aplodontia* and functional morphology in fossil aplodontines is needed to better understand patterns of digging mode evolution in this clade. Additionally, examination of cranial and postcranial functional indices informative for inference of chisel-tooth and head-lift digging may provide a more comprehensive picture of digging mode usage in *Aplodontia* and other cryptic extant species.

Using both phylogenetically naive and phylogenetically conditioned methods, the heteromyid *Chaetodipus intermedius* received greater mean WIS values for chisel-tooth digging than the three data-poor pocket gopher species. The sole other heteromyid species in the dataset, *Dipodomys microps*, was assigned a chisel-tooth ranking of 2; together with the paraphyletic recovery of Heteromyidae by Upham et al. (2019), these results might suggest that some degree of chisel-tooth digging may be plesiomorphic for Geomyoidea. The consistently low predicted chisel-tooth importance for the three data-poor pocket gopher species (*Orthogeomys grandis*, *Orthogeomys hispidus*, and *Zygogeomys trichopus*) suggests that chisel-tooth digging in *Cratogeomys* may be apomorphic rather than being the ancestral state for Geomyini and secondarily lost in *Geomys*. These patterns call into question what selective pressures and/or morphological or developmental constraints might drive frequent losses and secondary gains of incisor digging function within clades. In particular, shared ancestry with chisel-tooth digging taxa might confer a predisposition for re-evolving usage of this digging mode, especially with regard to phylogenetic inertia of morphologies associated with chisel-tooth digging (e.g., procumbent and deeply rooted incisors, tall occiputs).

The ultimate goal of many comparative studies with a predictive element is to apply the framework to the fossil record to reconstruct ecology, behavior, and function in extinct taxa. Cranial suture morphology presents an interesting challenge in this regard. Given the frequent correlations between suture complexity and function in the modern biota and the significance of sutures as an important taxonomically diagnostic tool in the vertebrate fossil record (Greenwood 1959, Roston et al. 2023, Kammerer et al. 2025), it seems logical to use suture morphology to illuminate patterns of function and ecology in extinct lineages (see Markey and Marshall 2007 for an application in early tetrapods). Nasofrontal suture complexity has been shown to correlate with inferred digging ability in dicynodont therapsids (Kammerer 2021), so the relationship between elevated craniofacial suture complexity and fossorial function is clearly not unique to Rodentia or even to Mammalia. However, the high degree of clade specificity in our findings and the substantial degree of intraspecific variation in suture complexity warrant caution when predicting digging mode importance or other aspects of function and ecology in fossils with this proxy. It will be crucial to consider evolutionary history and sample size (e.g., choosing well-represented species, comparing intraspecific variance in fossil taxa and related modern lineages) in future work on suture complexity in extinct lineages. For example, a certain complexity metric that predicts scratch digging importance in multiple rodent suborders may not translate well to other mammalian orders or non-mammalian synapsids, whereas a small sample size for a fossil taxon may not adequately capture intraspecific variance and could skew estimates of average suture complexity. The effects of taphonomic deformation, preservation, and even preparation should also be explored and accounted for when examining suture morphology in fossils. Deformation regime can affect shape analyses (e.g., geometric morphometrics) (Angielczyk and Sheets 2007, Kammerer et al. 2020, Ford et al. 2023, Hedrick 2023), and poor preparation, overpreparation, and weathering of the primary bone surface may contribute to skewed estimates of suture complexity (see Khurelbaatar et al. 2025 for a discussion of how suture morphology changes from the ectocranium to the endocranium). Our results underscore that a better understanding of the factors that influence suture morphology in modern animals and in the fossil record is needed before it can be considered a universally applicable proxy for digging mode and other aspects of functional ecology.

## Conclusions

Locomotor function is crucial to a species’ interactions with its environment. Even within a specific ecotype, locomotion is complex and multidimensional yet often oversimplified. Fossorial rodent species frequently utilize multiple distinct ways of digging, and such digging modes can be of varying importance, but this nuance is often neglected in comparative studies of morphology and function. Cranial suture morphology is relatively well studied in the context of cranial growth, diet, and static loading, but broader relationships between suture complexity, locomotion, and non-feeding ecology are more poorly understood. Using an ordinal ranking system for digging mode importance, cranial suture complexity data, including a new method of quantifying irregularity via spectral entropy, and Bayesian multilevel modeling, we demonstrate that aspects of suture phenotype can be used to predict fossorial function at the suborder level and above in Rodentia. Prior work has suggested that chisel-tooth and/or head-lift digging may correlate with elevated complexity in the premaxillofrontal and/or nasofrontal sutures. However, our results indicate that scratch digging most consistently correlates with increased complexity in the premaxillofrontal suture, whereas specialization for chisel-tooth digging correlates with decreased nasofrontal suture complexity. These findings highlight that locomotor ecology can be quantified in a nuanced, complex manner and that multiple axes of phenotypic variation within the same anatomical structure, such as a single cranial suture type, can reflect various aspects of function differently within and across clades.

## Acknowledgements

We are grateful to Anderson Feijo, Lauren Johnson, Adam Ferguson, Teresa Hsu, and Darrin Lunde for facilitating access to specimens and libraries. Many thanks to Jonathan Nations for sharing code and providing methodological feedback. Mark Webster, Graham Slater, Stephanie Smith, Danielle Adams, Callum Ross, Zhe-Xi Luo, Craig Byron, and Henry Fulghum provided helpful comments on the contents of the manuscript. This work was funded by the Geological Society of America Graduate Student Research Fund and the University of Chicago Hinds Fund.

