## Supplementary Table 1 for "Craniofacial suture complexity and digging mode importance in rodents"

| Species | Suborder | Family | ZygoConfig | Fossorial | Head-lift rank | Chisel-tooth rank | Scratch rank | Source(s) |
|---|---|---|---|---|---|---|---|---|
| Allactaga tetradactyla | Myomorpha | Dipodidae | Mymorphy | 1 | 2 | 2 | 3 | Eisenberg, John F. "The behavior patterns of desert rodents." Rodents in desert environments. Dordrecht: Springer Netherlands, 1975. 189-224. |
| Ammospermophilus leucurus | Sciuromorpha | Sciuridae | Sciuromorph | 1 | 1 | 1 | 3 | Lagaria, Anna, and Dionisios Youlatos. "Anatomical correlates to scratch digging in the forelimb of European ground squirrels (Spermophilus citellus)." Journal of Mammalogy 87.3 (2006): 563-570. |
| Aplodontia rufa | Sciuromorpha | Aplodontiidae | Protrogomorph | 1 | Unknown | Unknown | 3 | Anthony, Harold Elmer. "Habits of Aplodontia. Bulletin of the AMNH; v. 35, article 6." (1916). |
| Apomys insignis | Myomorpha | Muridae | Myomorph | 0 | 1 | 1 | 1 | Heaney, Lawrence R., et al. "Seven new species and a new subgenus of forest mice (Rodentia: Muridae: Apomys) from Luzon Island." Fieldiana Life and Earth Sciences 2011.2 (2011): 1-60. |
| Arborimus longicaudus | Myomorpha | Cricetidae | Myomorph | 1 | 1 | 1 | Unknown | Walker's Mammals of the World 6th Edition, Handbook of Mammals of the World |
| Arborimus pomo | Myomorpha | Cricetidae | Myomorph | 1 | 1 | 1 | Unknown | Walker's Mammals of the World 6th Edition, Handbook of Mammals of the World |
| Bandicota bengalensis | Myomorpha | Muridae | Myomorph | 1 | Unknown | 3 | Unknown | Kryštufek, Boris, et al. "Morphological evolution of the skull in closely related bandicoot rats: a comparative study using geometric morphometrics." Hystrix 27.2 (2016): 163. |
| Bandicota indica | Myomorpha | Muridae | Myomorph | 1 | 1 | 1 | 3 | Sridhara, Shakunthala. "Behavioural Ecology of Larger Bandicoot Rat, Bandicota indica (Bechstein), Burrow Ecology, Social and Feeding Behaviour." Animal Behavior in the Tropics: Vertebrates. Singapore: Springer Nature Singapore, 2025. 361-377. |
| Bathyergus suillus | Hystricomorpha | Bathyergidae | Protrogomorph* | 1 | 1 | 1 | 3 | Walker's Mammals of the World 6th Edition, Handbook of Mammals of the World |
| Callosciurus prevostii | Sciuromorpha | Sciuridae | Sciuromorph | 0 | 1 | 1 | 1 | Walker's Mammals of the World 6th Edition, Handbook of Mammals of the World |
| Cannomys badius | Myomorpha | Spalacidae | Myomorph | 1 | 1 | 3 | 2 | Walker's Mammals of the World 6th Edition, Handbook of Mammals of the World |
| Castor canadensis | Castorimorpha | Castoridae | Sciuromorph | 1 | 1 | 1 | 2 | Baker, B. W., and E. P. Hill. 2003. Beaver (Castor canadensis). Pages 288-310 in G. A. Feldhamer, B. C. Thompson, and J. A. Chapman, editors. Wild Mammals of North America: Biology, Management, and Conservation. Second Edition. The Johns Hopkins University Press, Baltimore, Maryland, USA. |
| Cavia tschudii | Hystricomorpha | Caviidae | Hystricomorph | 1 | Unknown | Unknown | Unknown | |
| Chaetodipus intermedius | Castorimorpha | Heteromyidae | Sciuromorph | Unknown | Unknown | Unknown | Unknown | |
| Chinchilla lanigera | Hystricomorpha | Chinchillidae | Hystricomorph | 0 | 1 | 1 | 1 | Walker's Mammals of the World 6th Edition, Handbook of Mammals of the World |
| Coendou rufescens | Hystricomorpha | Erethizontidae | Hystricomorph | 0 | 1 | 1 | 1 | Walker's Mammals of the World 6th Edition, Handbook of Mammals of the World |
| Cratogeomys castanops | Castorimorpha | Geomyidae | Sciuromorph | 1 | 1 | 2 | 3 | Walker's Mammals of the World 6th Edition, Handbook of Mammals of the World |
| Cratogeomys fumosus | Castorimorpha | Geomyidae | Sciuromorph | 1 | 1 | 2 | 3 | Walker's Mammals of the World 6th Edition, Handbook of Mammals of the World |
| Cryptomys hottentotus | Hystricomorpha | Bathyergidae | Protrogomorph* | 1 | 2 | 3 | 3 | Genelly, Richard E. "Ecology of the common mole-rat (Cryptomys hottentotus) in Rhodesia." Journal of Mammalogy 46.4 (1965): 647-665. |
| Ctenomys colburni | Hystricomorpha | Ctenomyidae | Hystricomorph | 1 | 1 | 3 | 3 | Walker's Mammals of the World 6th Edition, Handbook of Mammals of the World |
| Ctenomys dorsalis | Hystricomorpha | Ctenomyidae | Hystricomorph | 1 | 1 | 3 | 3 | Walker's Mammals of the World 6th Edition, Handbook of Mammals of the World |
| Ctenomys fulvus | Hystricomorpha | Ctenomyidae | Hystricomorph | 1 | 1 | 3 | 3 | Walker's Mammals of the World 6th Edition, Handbook of Mammals of the World |
| Ctenomys latro | Hystricomorpha | Ctenomyidae | Hystricomorph | 1 | 1 | 3 | 3 | Walker's Mammals of the World 6th Edition, Handbook of Mammals of the World |
| Ctenomys magellanicus | Hystricomorpha | Ctenomyidae | Hystricomorph | 1 | 1 | 3 | 3 | Walker's Mammals of the World 6th Edition, Handbook of Mammals of the World |
| Ctenomys mendocinus | Hystricomorpha | Ctenomyidae | Hystricomorph | 1 | 1 | 2 | 3 | Camin, S., L. Madoery, and V. Roig. "The burrowing behavior of Ctenomys mendocinus (Rodentia)." (1995): 9-18. |
| Ctenomys opimus | Hystricomorpha | Ctenomyidae | Hystricomorph | 1 | 1 | 1 | 3 | Mammals of the Neotropics Vol. 2 |
| Ctenomys sericeus | Hystricomorpha | Ctenomyidae | Hystricomorph | 1 | 1 | 3 | 3 | Walker's Mammals of the World 6th Edition, Handbook of Mammals of the World |
| Ctenomys talarum | Hystricomorpha | Ctenomyidae | Hystricomorph | 1 | 1 | 3 | 3 | Walker's Mammals of the World 6th Edition, Vassallo, Aldo I. "Functional morphology, comparative behaviour, and adaptation in two sympatric subterranean rodents genus Ctenomys (Caviomorpha: Octodontidae)." Journal of Zoology 244.3 (1998): 415-427. |
| Ctenomys tuconax | Hystricomorpha | Ctenomyidae | Hystricomorph | 1 | 1 | 3 | 3 | Walker's Mammals of the World 6th Edition, Handbook of Mammals of the World |
| Cuniculus paca | Hystricomorpha | Cuniculidae | Hystricomorph | 1 | 1 | 1 | 2 | Walker's Mammals of the World 6th Edition, Handbook of Mammals of the World |
| Cynomys gunnisoni | Sciuromorpha | Sciuridae | Sciuromorph | 1 | 1 | 1 | 3 | Scheetz, Luke. Scratching Beneath the Surface: Quantifications of Muscle Architecture and Myosin Heavy Chain Content in the Forelimbs of Black-tailed Prairie Dogs (Cynomys, Rodentia). MS thesis. Youngstown State University, 2024. |
| Cynomys leucurus | Sciuromorpha | Sciuridae | Sciuromorph | 1 | 1 | 1 | 3 | Scheetz, Luke. Scratching Beneath the Surface: Quantifications of Muscle Architecture and Myosin Heavy Chain Content in the Forelimbs of Black-tailed Prairie Dogs (Cynomys, Rodentia). MS thesis. Youngstown State University, 2024. |
| Cynomys ludovicianus | Sciuromorpha | Sciuridae | Sciuromorph | 1 | 1 | 1 | 3 | Scheetz, Luke. Scratching Beneath the Surface: Quantifications of Muscle Architecture and Myosin Heavy Chain Content in the Forelimbs of Black-tailed Prairie Dogs (Cynomys, Rodentia). MS thesis. Youngstown State University, 2024. |
| Desmodillus auricularis | Myomorpha | Muridae | Myomorph | 1 | 1 | 1 | 3 | Keogh, H. J. "Behaviour and breeding in captivity of the Namaqua gerbil Desmodillus auricularis (Cricetidae: Gerbillinae)." African Zoology 8.2 (1973): 231-239. |
| Dinomys branickii | Hystricomorpha | Dinomyidae | Hystricomorph | 1 | 1 | 1 | 2 | Walker's Mammals of the World 6th Edition, Handbook of Mammals of the World |
| Dipodomys microps | Castorimorpha | Heteromyidae | Sciuromorph | 1 | 1 | 2 | 3 | Eisenberg, John F. "The behavior patterns of desert rodents." Rodents in desert environments. Dordrecht: Springer Netherlands, 1975. 189-224. |
| Ellobius fuscocapillus | Myomorpha | Cricetidae | Myomorph | 1 | 1 | 3 | 2 | Walker's Mammals of the World 6th Edition, Handbook of Mammals of the World |
| Ellobius talpinus | Myomorpha | Cricetidae | Myomorph | 1 | 1 | 3 | 2 | Walker's Mammals of the World 6th Edition, Handbook of Mammals of the World |
| Eospalax fontanierii | Myomorpha | Spalacidae | Myomorph | 1 | 2 | 2 | 3 | Walker's Mammals of the World 6th Edition, Handbook of the Mammals of the World |
| Eospalax smithii | Myomorpha | Spalacidae | Myomorph | 1 | 2 | 2 | 3 | Walker's Mammals of the World 6th Edition, Handbook of Mammals of the World |
| Erethizon dorsatum | Hystricomorpha | Erethizontidae | Hystricomorph | 0 | 1 | 1 | 1 | Johnson, Judith. "Biology of the Porcupine (Erethizon dorsatum) in northwestern South Dakota." (1977). |
| Fukomys amatus | Hystricomorpha | Bathyergidae | Protrogomorph* | 1 | 1 | 3 | 3 | Walker's Mammals of the World 6th Edition, Handbook of Mammals of the World |
| Fukomys darlingi | Hystricomorpha | Bathyergidae | Protrogomorph* | 1 | 1 | 3 | 3 | Walker's Mammals of the World 6th Edition, Handbook of Mammals of the World |
| Fukomys ochraceocinereus | Hystricomorpha | Bathyergidae | Protrogomorph* | 1 | 1 | 3 | 3 | Walker's Mammals of the World 6th Edition, Handbook of Mammals of the World |
| Fukomys zechi | Hystricomorpha | Bathyergidae | Protrogomorph* | 1 | 1 | 3 | 3 | Walker's Mammals of the World 6th Edition, Handbook of Mammals of the World |
| Geomys attwateri | Castorimorpha | Geomyidae | Sciuromorph | 1 | 1 | 1 | 3 | Walker's Mammals of the World 6th Edition, Handbook of Mammals of the World |
| Geomys bursarius | Castorimorpha | Geomyidae | Sciuromorph | 1 | 1 | 1 | 3 | Walker's Mammals of the World 6th Edition, Handbook of Mammals of the World |
| Geomys personatus | Castorimorpha | Geomyidae | Sciuromorph | 1 | 1 | 1 | 3 | Walker's Mammals of the World 6th Edition, Handbook of Mammals of the World |
| Geomys pinetis | Castorimorpha | Geomyidae | Sciuromorph | 1 | 1 | 1 | 3 | Walker's Mammals of the World 6th Edition, Handbook of Mammals of the World |
| Georychus capensis | Hystricomorpha | Bathyergidae | Protrogomorph* | 1 | 1 | 3 | 2 | Walker's Mammals of the World 6th Edition, Handbook of Mammals of the World |
| Gerbillus gerbillus | Myomorpha | Muridae | Myomorph | 1 | 1 | 1 | 3 | Agrawal, V. C. "Skull adaptations in fossorial rodents." (1967): 300-312, Walker's Mammals of the World 6th Edition |
| Glaucomys sabrinus | Sciuromorpha | Sciuridae | Sciuromorph | 0 | 1 | 1 | 1 | Walker's Mammals of the World 6th Edition, Handbook of Mammals of the World |
| Glaucomys volans | Sciuromorpha | Sciuridae | Sciuromorph | 0 | 1 | 1 | 1 | Walker's Mammals of the World 6th Edition, Handbook of Mammals of the World |
| Glis glis | Sciuromorpha | Gliridae | Myomorph* | 0 | 1 | 1 | 1 | Walker's Mammals of the World 6th Edition, Handbook of Mammals of the World |
| Graphiurus murinus | Sciuromorpha | Gliridae | Myomorph* | 0 | 1 | 1 | 1 | Perrin, M. R. "Notes on the activity patterns of 12 species of southern African rodents and a new design of activity monitor." African Zoology 16.4 (1981): 248-258. |
| Heliophobius argenteocinereus | Hystricomorpha | Bathyergidae | Protrogomorph* | 1 | 2 | 3 | 2 | Walker's Mammals of the World 6th Edition, Handbook of Mammals of the World |
| Heliosciurus rufobrachium | Sciuromorpha | Sciuridae | Sciuromorph | 0 | 1 | 1 | 1 | Walker's Mammals of the World 6th Edition, Handbook of Mammals of the World |
| Heterocephalus glaber | Hystricomorpha | Heterocephalidae | Protrogomorph* | 1 | 2 | 3 | 2 | Tucker, Richard. "The digging behavior and skin differentiations in Heterocephalus glaber." Journal of Morphology 168.1 (1981): 51-71. |
| Hydrochoerus hydrochaeris | Hystricomorpha | Hydrochoeridae | Hystricomorph | 0 | 1 | 1 | 1 | Walker's Mammals of the World 6th Edition, Handbook of Mammals of the World |
| Hystrix crassispinis | Hystricomorpha | Hystricidae | Hystricomorph | 1 | 1 | 1 | 2 | Walker's Mammals of the World 6th Edition, Handbook of Mammals of the World |
| Isothrix bistriata | Hystricomorpha | Echimyidae | Hystricomorph | 0 | 1 | 1 | 1 | Walker's Mammals of the World 6th Edition, Handbook of Mammals of the World |
| Jaculus jaculus | Myomorpha | Dipodidae | Myomorph | 1 | 2 | 3 | 3 | Eisenberg, John F. "The behavior patterns of desert rodents." Rodents in desert environments. Dordrecht: Springer Netherlands, 1975. 189-224. |
| Lagidium viscacia | Hystricomorpha | Chinchillidae | Hystricomorph | 0 | 1 | 1 | 1 | Walker's Mammals of the World 6th Edition, Handbook of Mammals of the World |
| Lagostomus maximus | Hystricomorpha | Chinchillidae | Hystricomorph | 1 | 2 | 1 | 3 | Walker's Mammals of the World 6th Edition, Handbook of Mammals of the World |
| Marmota monax | Sciuromorpha | Sciuridae | Sciuromorph | 1 | 2 | 1 | 3 | Kwiecinski, Gary G. "Marmota monax." Mammalian Species 591 (1998): 1-8. |
| Microtus californicus | Myomorpha | Cricetidae | Myomorph | Unknown | Unknown | Unknown | Unknown | |
| Microtus pennsylvanicus | Myomorpha | Cricetidae | Myomorph | 1 | 1 | 1 | 2 | Webster, Daniel G., et al. "Digging behavior in 12 taxa of muroid rodents." Animal Learning & Behavior 9.2 (1981): 173-177. |
| Mus musculus | Myomorpha | Muridae | Myomorph | 1 | 1 | 1 | 3 | Webster, Daniel G., et al. "Digging behavior in 12 taxa of muroid rodents." Animal Learning & Behavior 9.2 (1981): 173-177. |
| Myocastor coypus | Hystricomorpha | Echimyidae | Hystricomorph | 1 | 1 | 1 | 2 | Woods, Charles A., et al. "Myocastor coypus." Mammalian species 398 (1992): 1-8. |
| Myospalax psilurus | Myomorpha | Spalacidae | Myomorph | 1 | 2 | 2 | 3 | Walker's Mammals of the World 6th Edition, Handbook of Mammals of the World |
| Myospalax myospalax | Myomorpha | Spalacidae | Myomorph | 1 | 2 | 2 | 3 | Walker's Mammals of the World 6th Edition, Handbook of Mammals of the World |
| Neotoma fuscipes | Myomorpha | Cricetidae | Myomorph | 0 | 1 | 1 | 1 | Walker's Mammals of the World 6th Edition, Handbook of Mammals of the World |
| Nesokia indica | Myomorpha | Muridae | Myomorph | 1 | Unknown | 3 | Unknown | Kryštufek, Boris, et al. "Morphological evolution of the skull in closely related bandicoot rats: a comparative study using geometric morphometrics." Hystrix 27.2 (2016): 163. |
| Notomys alexis | Myomorpha | Muridae | Myomorph | 1 | 2 | 1 | 3 | Stanley, Meredith. "An ethogram of the hopping mouse, Notomys alexis." Zeitschrift für Tierpsychologie 29.3 (1971): 225-258. |
| Octodon degus | Hystricomorpha | Octodontidae | Hystricomorph | 1 | 2 | 2 | 3 | Ebensperger, Luis A., and Francisco Bozinovic. "Energetics and burrowing behaviour in the semifossorial degu Octodon degus (Rodentia: Octodontidae)." Journal of Zoology 252.2 (2000): 179-186. |
| Octodontomys gliroides | Hystricomorpha | Octodontidae | Hystricomorph | 1 | Unknown | Unknown | Unknown | |
| Ondatra zibethicus | Myomorpha | Cricetidae | Myomorph | 1 | 1 | 1 | 2 | Walker's Mammals of the World 6th Edition, Handbook of Mammals of the World |
| Orthogeomys grandis | Castorimorpha | Geomyidae | Sciuromorph | 1 | Unknown | Unknown | Unknown | |
| Orthogeomys hispidus | Castorimorpha | Geomyidae | Sciuromorph | 1 | Unknown | Unknown | Unknown | |
| Pedetes capensis | Anomaluromorph | Pedetidae | Hystricomorph | 1 | 1 | 2 | 3 | Butynski, Thomas M., and Rosanna Mattingly. "Burrow structure and fossorial ecology of the springhare Pedetes capensis in Botswana." African Journal of Ecology 17.4 (1979): 205-215. |
| Peromyscus californicus | Myomorpha | Cricetidae | Myomorph | 0 | 1 | 1 | 1 | Merritt, Joseph F. "Peromyscus californicus." Mammalian Species 85 (1978): 1-6. |
| Peromyscus maniculatus | Myomorpha | Cricetidae | Myomorph | 1 | 1 | 1 | 3 | Webster, Daniel G., et al. "Digging behavior in 12 taxa of muroid rodents." Animal Learning & Behavior 9.2 (1981): 173-177. |
| Petaurista elegans | Sciuromorpha | Sciuridae | Sciuromorph | 0 | 1 | 2 | 2 | Walker's Mammals of the World 6th Edition, Handbook of Mammals of the World |
| Phloeomys pallidus | Myomorpha | Muridae | Myomorph | 0 | 1 | 1 | 1 | Walker's Mammals of the World 6th Edition, Handbook of Mammals of the World |
| Proechimys brevicauda | Hystricomorpha | Echimyidae | Hystricomorph | 0 | 1 | 1 | 1 | Walker's Mammals of the World 6th Edition, Handbook of Mammals of the World |
| Pseudomys nanus | Myomorpha | Muridae | Myomorph | 0 | 1 | 1 | 1 | Walker's Mammals of the World 6th Edition, Handbook of Mammals of the World |
| Rattus exulans | Myomorpha | Muridae | Myomorph | 0 | 1 | 1 | 1 | Walker's Mammals of the World 6th Edition, Handbook of Mammals of the World |
| Rattus norvegicus | Myomorpha | Muridae | Myomorph | 1 | 1 | 2 | 3 | Pisano, Rocci G., and Tracy I. Storer. "Burrows and feeding of the Norway rat." Journal of Mammalogy 29.4 (1948): 374-383. |
| Ratufa indica | Sciuromorpha | Sciuridae | Sciuromorph | 0 | 1 | 1 | 1 | Walker's Mammals of the World 6th Edition, Handbook of Mammals of the World |
| Rhizomys pruinosus | Myomorpha | Spalacidae | Myomorph | 1 | 1 | 3 | 2 | Walker's Mammals of the World 6th Edition, Handbook of Mammals of the World |
| Rhizomys sinensis | Myomorpha | Spalacidae | Myomorph | 1 | 1 | 3 | 2 | Walker's Mammals of the World 6th Edition, Handbook of Mammals of the World |
| Rhizomys sumatrensis | Myomorpha | Spalacidae | Myomorph | 1 | 1 | 3 | 2 | Walker's Mammals of the World 6th Edition, Handbook of Mammals of the World |
| Sciurillus pusillus | Sciuromorpha | Sciuridae | Sciuromorph | 0 | 1 | 1 | 1 | Youlatos, Dionisios. "Substrate use and locomotor modes of the Neotropical pygmy squirrel Sciurillus pusillus (E. Geoffroy, 1803) in French Guyana." Zoological Studies 50.6 (2011): 745-750. |

| **Species** | **Suborder** | **Family** | **ZygoConfig** | Fossorial | Head-lift rank | Chisel-tooth rank | Scratch rank | **Source(s)** |
|---|---|---|---|---|---|---|---|---|
| Sciurus aberti | Sciuromorpha | Sciuridae | Sciuromorph | 0 | 1 | 1 | 1 | Walker's Mammals of the World 6th Edition, Handbook of Mammals of the World |
| Spalacopus cyanus | Hystricomorpha | Octodontidae | Hystricomorph | 1 | 1 | 3 | 3 | Walker's Mammals of the World 6th Edition, Handbook of Mammals of the World |
| Spalax leucodon | Myomorpha | Spalacidae | Myomorph | 1 | 3 | 3 | 2 | Walker's Mammals of the World 6th Edition, Handbook of Mammals of the World |
| Spermophilus citellus | Sciuromorpha | Sciuridae | Sciuromorph | 1 | 1 | 2 | 3 | Ramos-Lara, Nicolás, et al. "Spermophilus citellus (Rodentia: sciuridae)." Mammalian Species 46.913 (2014): 71-87. |
| Tachyoryctes splendens | Myomorpha | Spalacidae | Myomorph | 1 | 2 | 3 | 2 | Walker's Mammals of the World 6th Edition, Handbook of Mammals of the World |
| Tamias striatus | Sciuromorpha | Sciuridae | Sciuromorph | 1 | 1 | 1 | 3 | Yahner, Richard H. "The adaptive nature of the social system and behavior in the eastern chipmunk, Tamias striatus." Behavioral Ecology and Sociobiology 3.4 (1978): 397-427. |
| Thomomys bottae | Castorimorpha | Geomyidae | Sciuromorph | 1 | 1 | 2 | 3 | Walker's Mammals of the World 6th Edition, Marcy, Ariel E., et al. "Morphological adaptations for digging and climate-impacted soil properties define pocket gopher (Thomomys spp.) distributions." PLoS One 8.5 (2013): e64935. |
| Thomomys bulbivorus | Castorimorpha | Geomyidae | Sciuromorph | 1 | 1 | 2 | 3 | Walker's Mammals of the World 6th Edition, Marcy, Ariel E., et al. "Morphological adaptations for digging and climate-impacted soil properties define pocket gopher (Thomomys spp.) distributions." PLoS One 8.5 (2013): e64935. |
| Thomomys monticola | Castorimorpha | Geomyidae | Sciuromorph | 1 | 1 | 3 | 3 | Walker's Mammals of the World 6th Edition, Marcy, Ariel E., et al. "Morphological adaptations for digging and climate-impacted soil properties define pocket gopher (Thomomys spp.) distributions." PLoS One 8.5 (2013): e64935. |
| Thomomys talpoides | Castorimorpha | Geomyidae | Sciuromorph | 1 | 1 | 3 | 3 | Walker's Mammals of the World 6th Edition, Marcy, Ariel E., et al. "Morphological adaptations for digging and climate-impacted soil properties define pocket gopher (Thomomys spp.) distributions." PLoS One 8.5 (2013): e64935. |
| Thryonomys gregorianus | Hystricomorpha | Thryonomyidae | Hystricomorph | 1 | 1 | 1 | 2 | Walker's Mammals of the World 6th Edition, Handbook of Mammals of the World |
| Thryonomys swinderianus | Hystricomorpha | Thryonomyidae | Hystricomorph | 1 | 1 | 1 | 2 | Walker's Mammals of the World 6th Edition, Handbook of Mammals of the World |
| Xerus inauris | Sciuromorpha | Sciuridae | Sciuromorph | 1 | 1 | 2 | 3 | Herzig-Straschil, Barbara. "On the biology of Xerus inauris (Zimmermann, 1780)(Rodentia, Sciuridae)." Zeitschrift für Säugetierkunde 43 (1978): 262-278. |
| Zygogeomys trichopus | Castorimorpha | Geomyidae | Sciuromorphy | 1 | Unknown | Unknown | Unknown | |
