## Supplementary Table 1 for "Craniofacial suture complexity and digging mode importance in rodents"

**Supplementary Figure and Table Captions**

Fig. 1: Procrustes-superimposed sutures before (blue) and after (orange) sliding of the semilandmarks, demonstrating minimal distortion (left), moderate distortion (center), and high distortion (right) of curve shape caused by the sliding process. Museum abbreviations: Field Museum of Natural History (FMNH), Smithsonian National Museum of Natural History (NMNH).

Fig. 2: Correlation plot of Spearman’s rank correlations and associated p-values between the power spectrum density (PSD) and spectral entropy (SE) metrics derived from the non-slid and slid semilandmark datasets for each suture. Despite sliding sometimes resulting in shape differences (see Fig. 1), the sliding process has minimal effect on calculated PSD and spectral entropy value. Suture abbreviations: premaxillofrontal (PMF), nasofrontal (NF). P-value abbreviations: * = p < 0.05, ** = p < 0.01, *** = p < 0.001.

Fig. 3: Intercept-only prior predictive checks for each digging mode based on the known distribution of ranks among data-rich species. *y* represents the known distribution of ranks for each digging mode, and *y*rep represents the replicated data (500 draws) using the intercept priors. Intercept priors for each digging mode include: scratch = normal(-0.75, 1), chisel-tooth = normal(0.5, 0.75), head-lift = normal(2.5, 1.5).

Fig. 4: Posterior predictive checks using 500 randomly sampled posterior draws for each model with scratch digging rank as the response. From top left: premaxillofrontal sinuosity index (SI), nasofrontal SI, premaxillofrontal power spectrum density (PSD), nasofrontal PSD, premaxillofrontal spectral entropy, nasofrontal spectral entropy. *y* represents the known distribution of ranks for each digging mode, and *y*rep is the replicated data.

Fig. 5: Posterior predictive checks using 500 randomly sampled posterior draws for each model with chisel-tooth digging rank as the response. From top left: premaxillofrontal sinuosity index (SI), nasofrontal SI, premaxillofrontal power spectrum density (PSD), nasofrontal PSD, premaxillofrontal spectral entropy, nasofrontal spectral entropy. *y* represents the known distribution of ranks for each digging mode, and *y*rep is the replicated data.

Fig. 6: Posterior predictive checks using 500 randomly sampled posterior draws for each model with head-lift digging rank as the response. From top left: premaxillofrontal sinuosity index (SI), nasofrontal SI, premaxillofrontal power spectrum density (PSD), nasofrontal PSD, premaxillofrontal spectral entropy, nasofrontal spectral entropy. *y* represents the known distribution of ranks for each digging mode, and *y*rep is the replicated data.

Fig. 7: A: 89% credible intervals (CIs) of the posterior conditional effect sizes of each suture complexity metric on digging mode importance rank for the entire dataset (“grand mean”) and each suborder for the models discussed in the main text, as well as models with scratch rank as the response and binary chisel-tooth and head-lift usage as covariates and models with chisel-tooth rank as the response and binary scratch and head-lift usage as covariates. An effect is considered significant if its 89% CI excludes 0. Suture complexity metrics are listed on the left side of the y-axis, and data subsets (i.e., grand mean and each suborder) are listed on the right side of the y-axis. Teal hues represent models with significant negative effects, and orange hues represent models with significant positive effects. Suture abbreviations: premaxillofrontal (PMF), nasofrontal (NF).

Table 1: Taxonomic and ecological/behavioral data, along with the main sources used to assign rankings, for each species. Fossoriality is coded as a binary variable, with 0 indicating a non-fossorial lifestyle and 1 indicating digging or otherwise moving earthbound substrate as a regular behavioral occurrence. Digging mode rankings range from 1 (the digging mode is extremely rarely or never used) to 3 (the digging mode is crucial to the species’ overall lifestyle); see Methods for more detailed description of the ranking system.
