## Supplementary figures and images for "Craniofacial suture complexity and digging mode importance in rodents"

### Supplementary Fig. 1

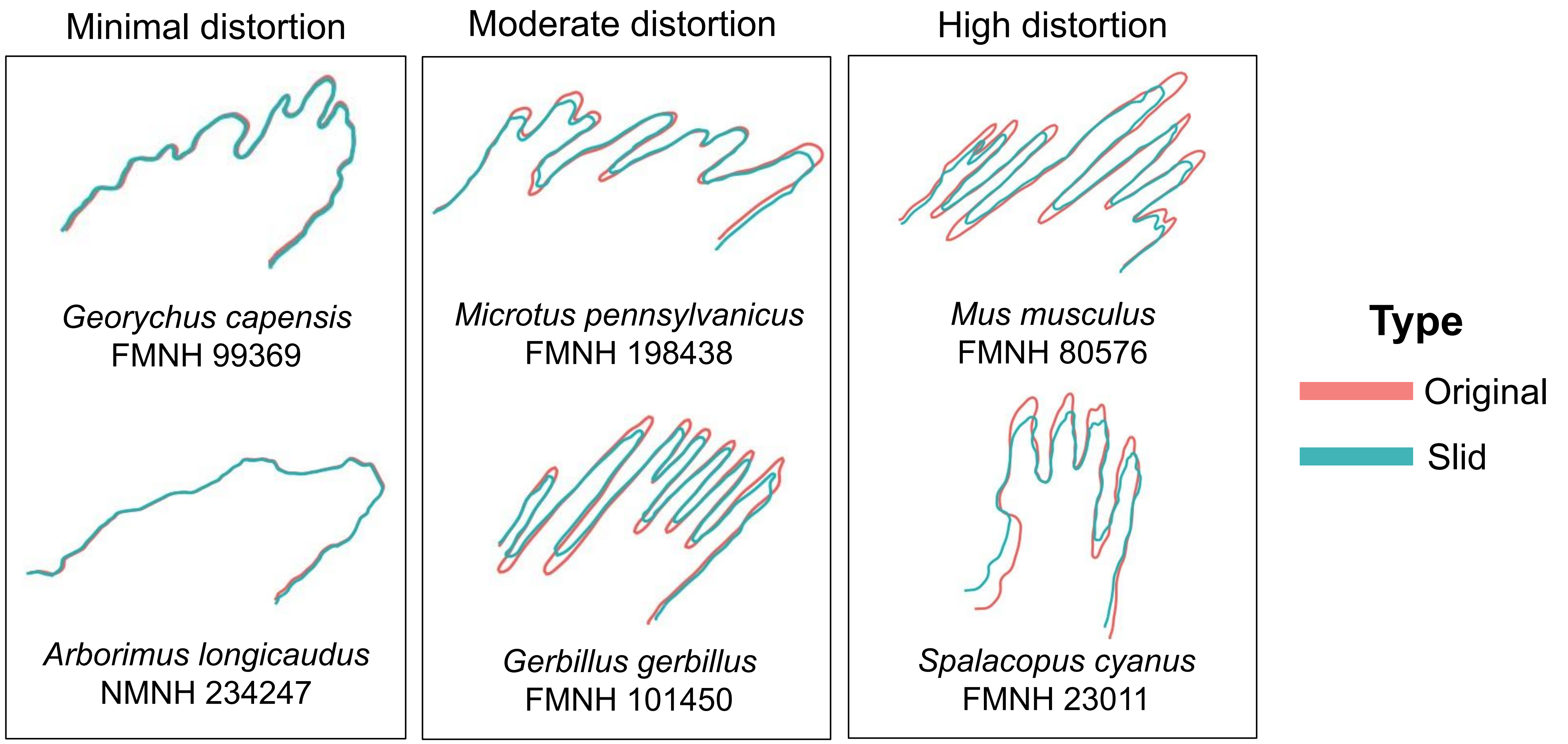

### Supplementary Fig. 2

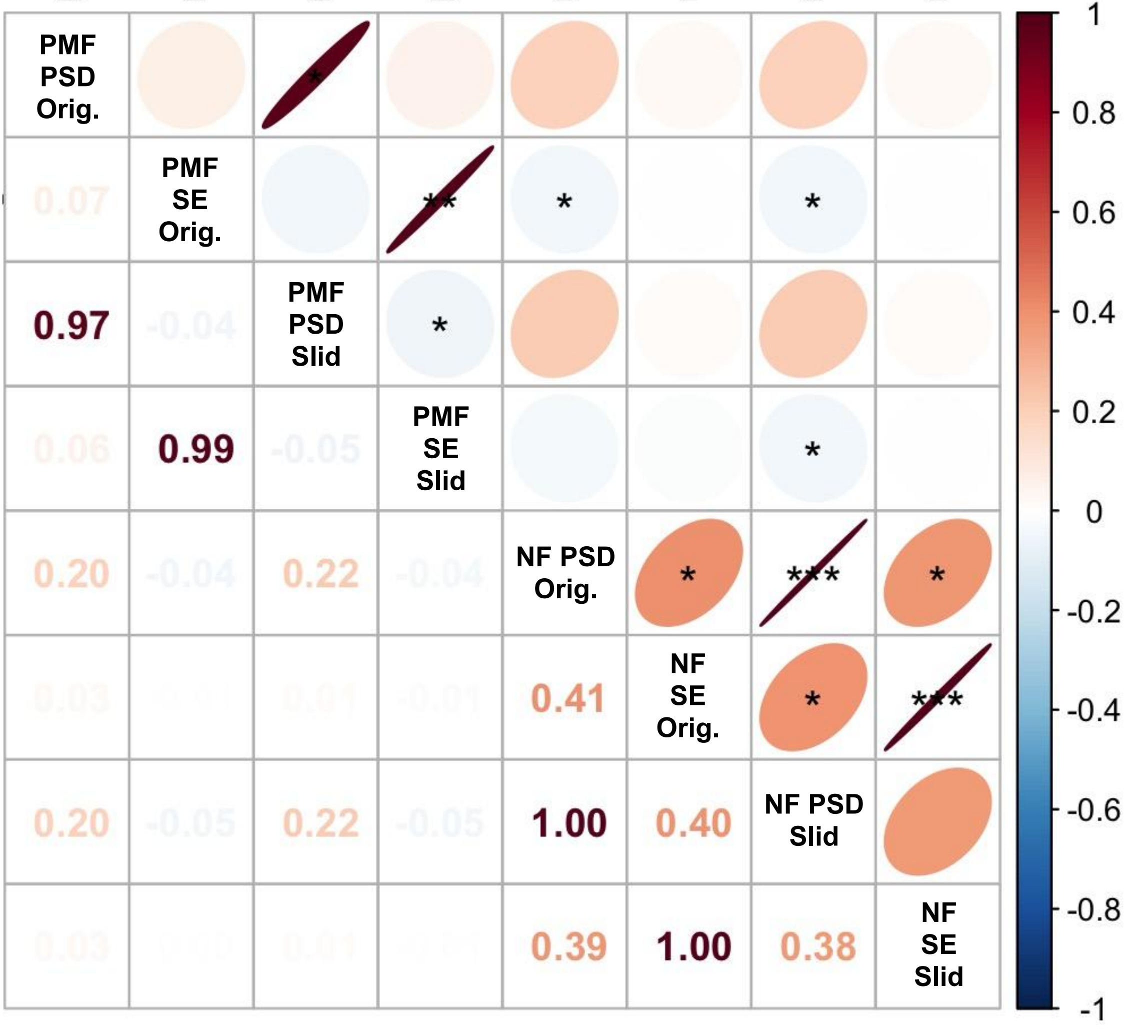

### Supplementary Fig. 3

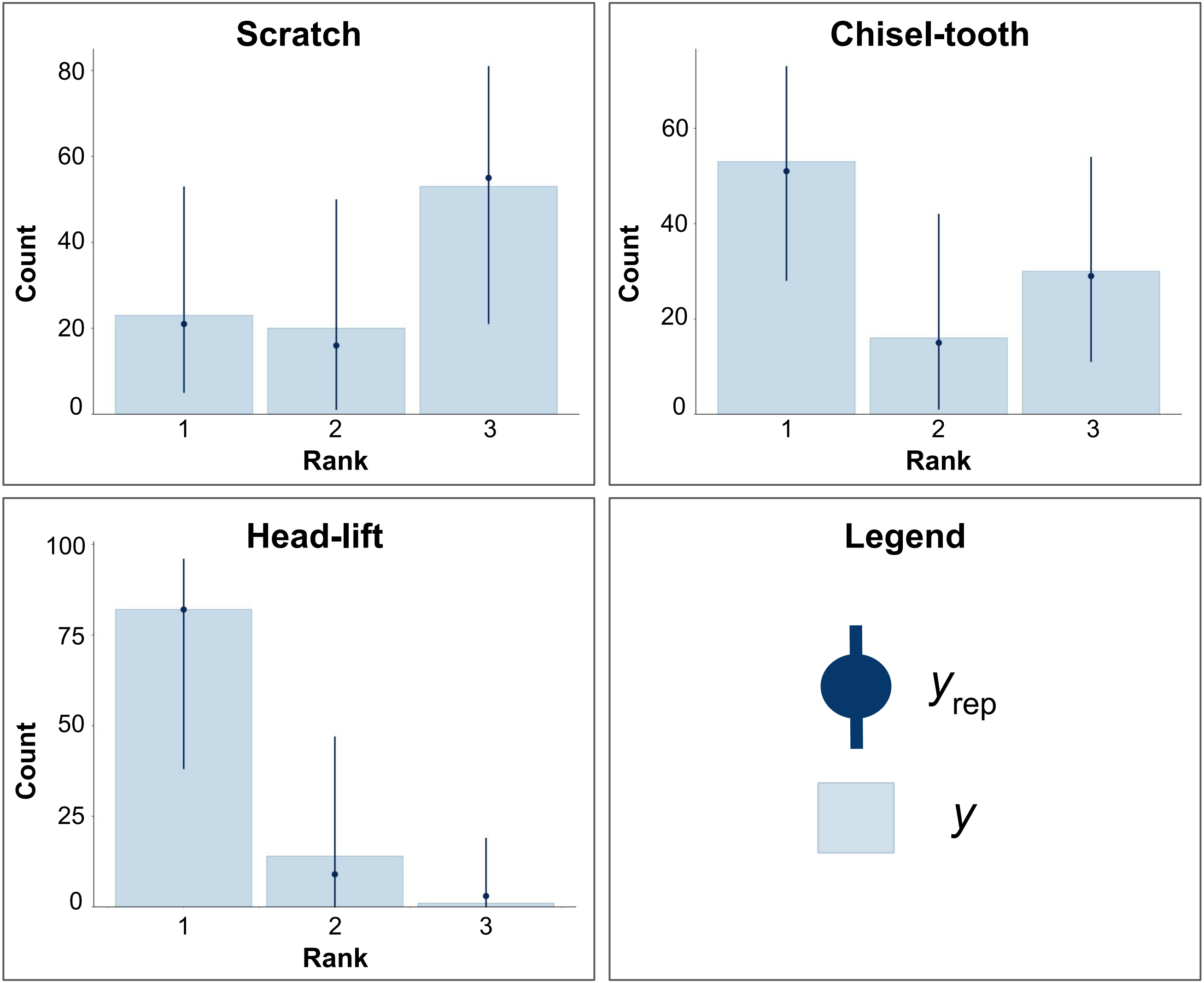

### Supplementary Fig. 7

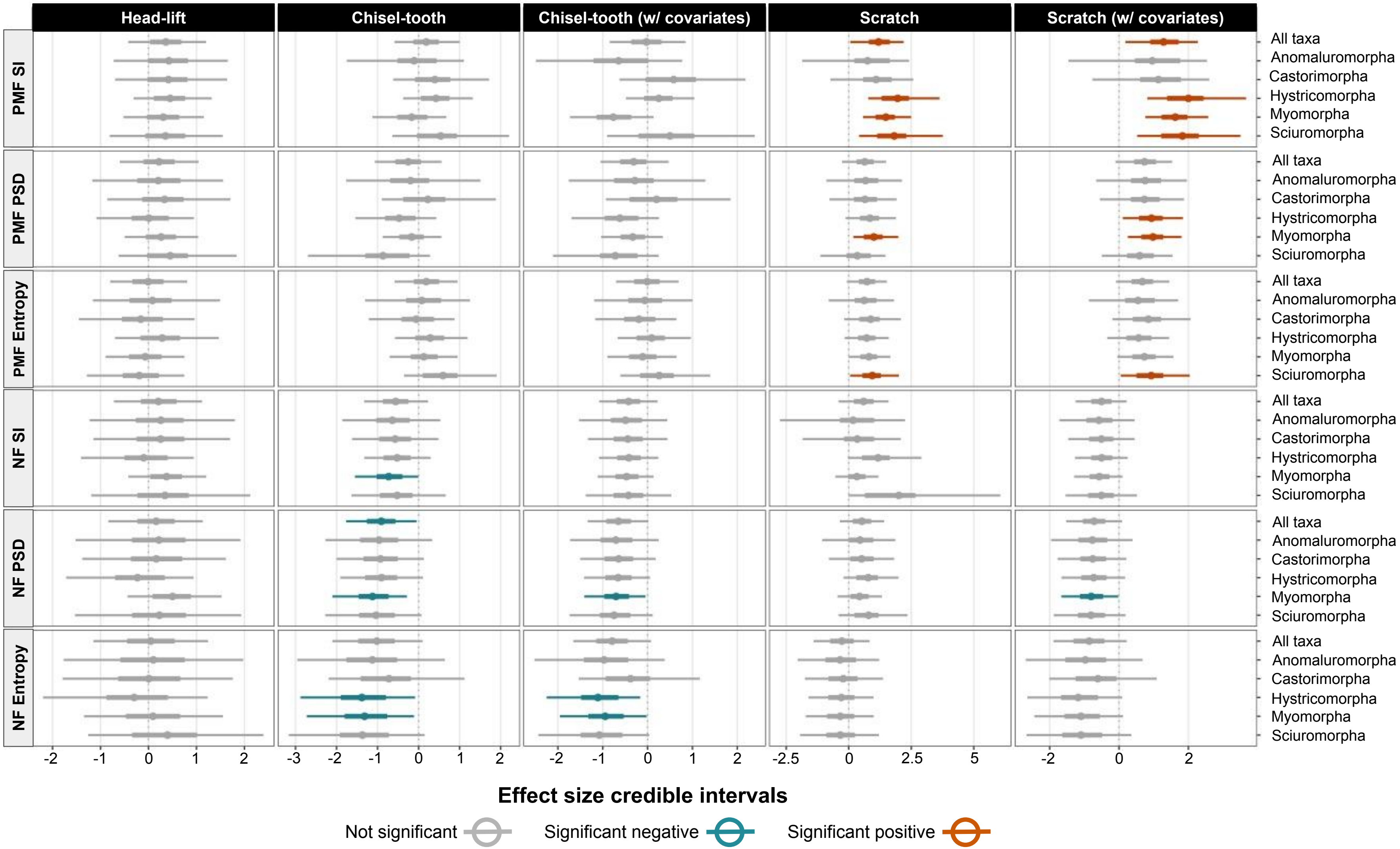

### Supplementary Figs. 4-6

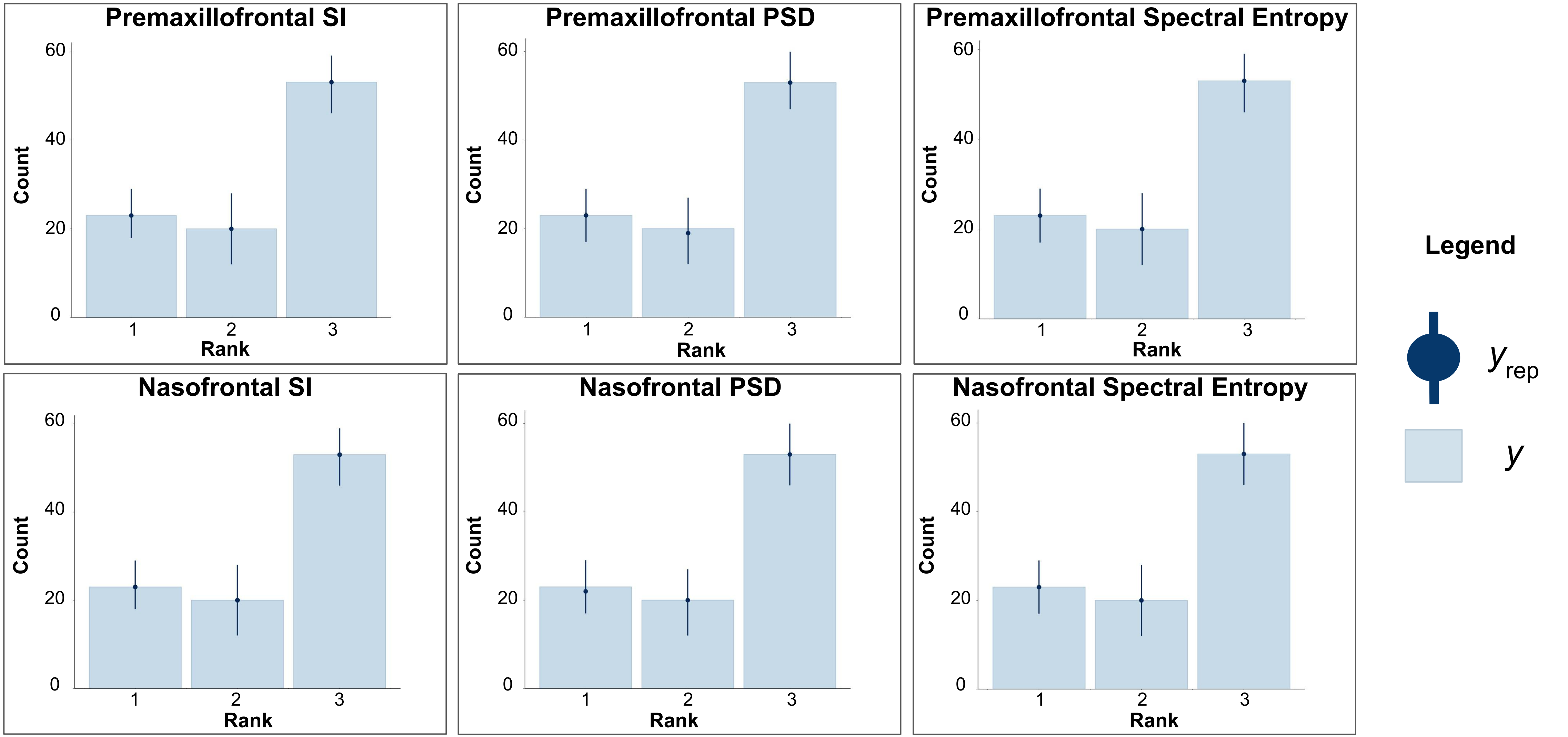

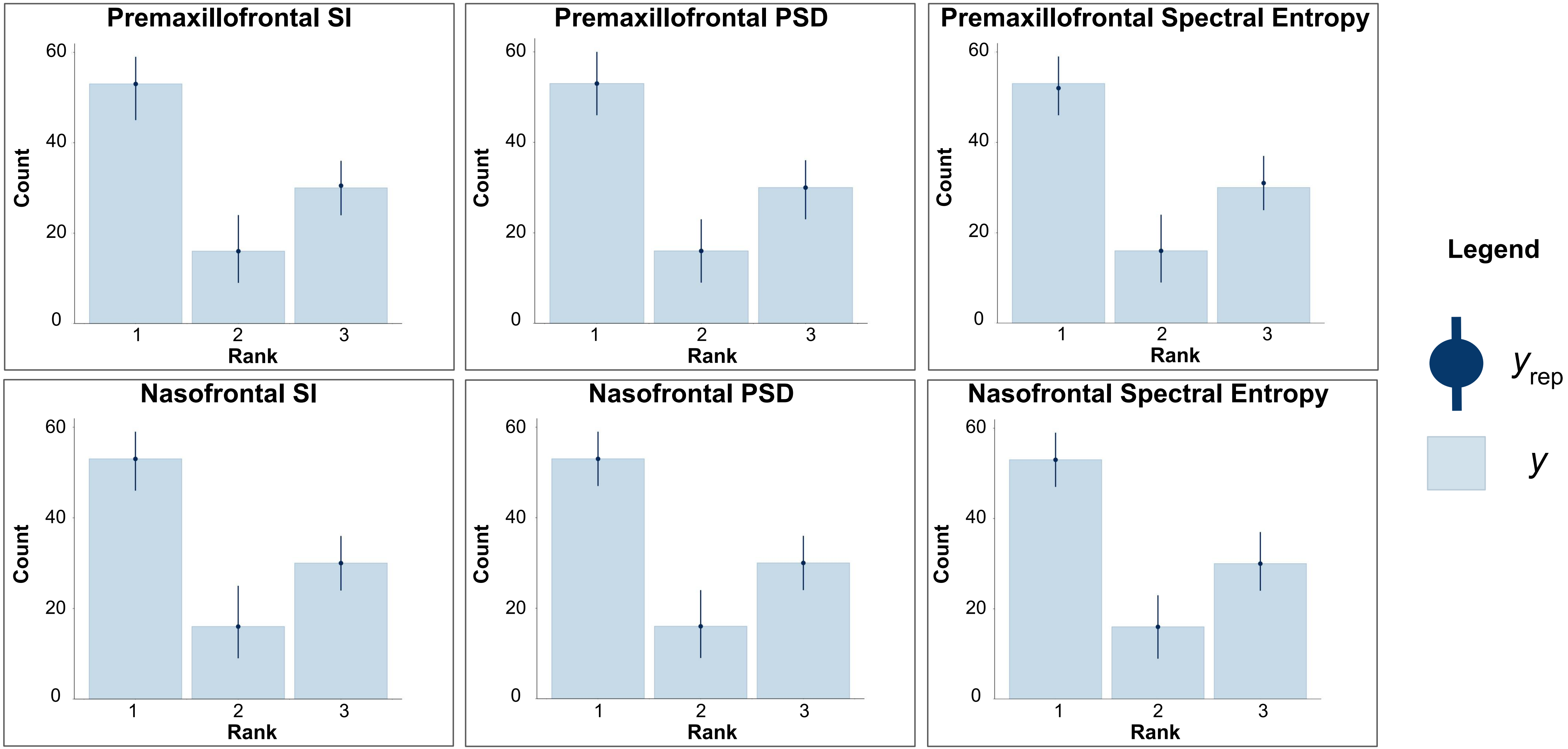

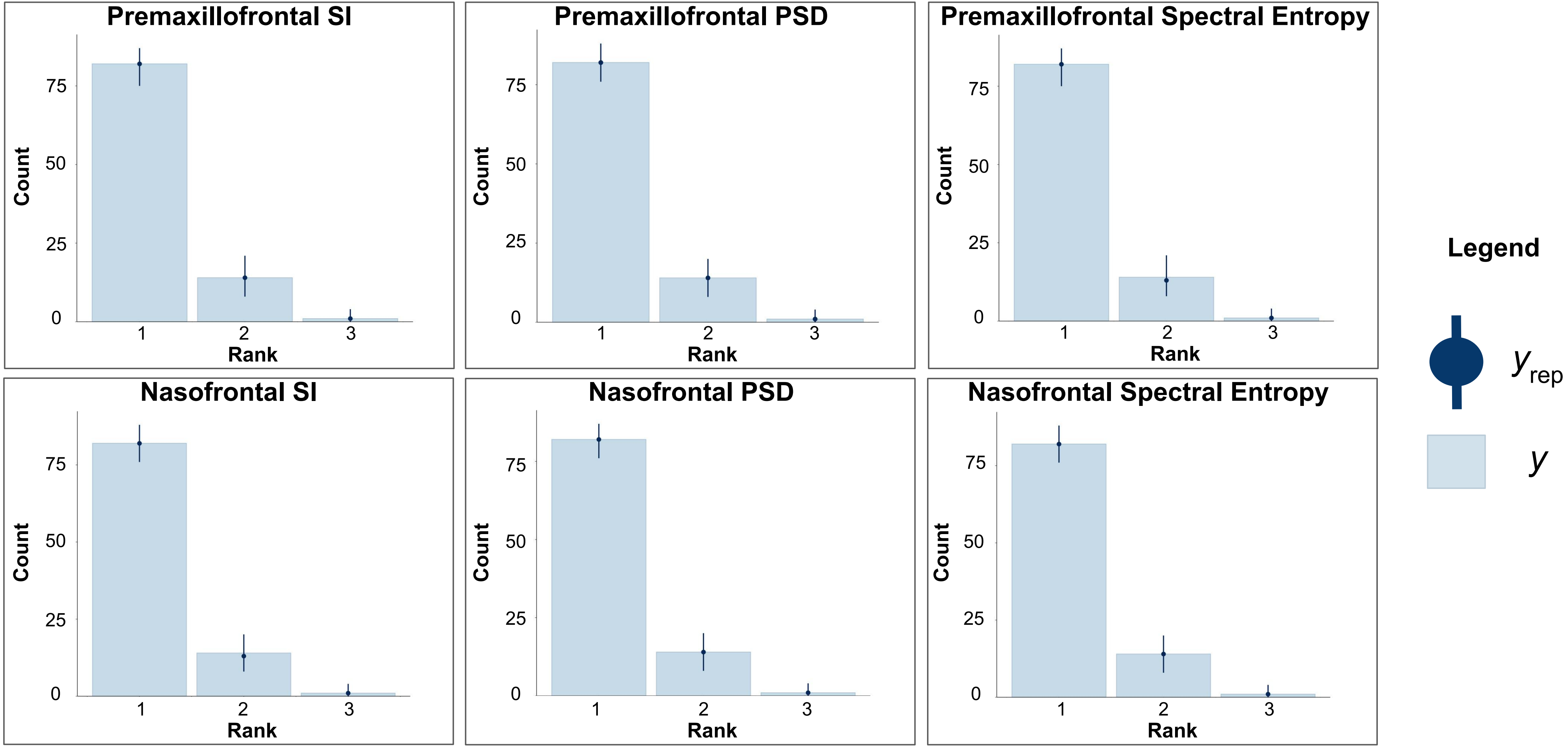
